# Neurotrophic and immunomodulatory lanostane triterpenoids from wood-inhabiting Basidiomycota

**DOI:** 10.1101/2022.10.27.513914

**Authors:** Khadija Hassan, Blondelle Matio Kemkuignou, Marco Kirchenwitz, Kathrin Wittstein, Monique Rascher, Clara Chepkirui, Josphat C. Matasyoh, Cony Decock, Reinhard Köster, Theresia E. B. Stradal, Marc Stadler

## Abstract

Neurotrophins such as nerve growth factor (*ngf*) and brain derived neurotrophic factor (*bdnf*) play important roles in the central nervous system. They are potential therapeutic drugs for treatment of neurodegerative diseases including Alzheimer’s disease and Parkinson’s disease. In this study, we investigated the neurotrophic properties of triterpenes isolated from fruiting bodies of *Laetiporus sulphureus* and a mycelial culture of *Antrodia* sp. MUCL 56049. The structures of the isolated compounds were elucidated based on nuclear magnetic resonance (NMR) spectroscopy in combination with high-resolution Electrospray mass spectrometry (HR-ESIMS). The secondary metabolites were tested for neurotrophin (*ngf* and *bdnf*) expression levels on human astrocytoma 1321N1 cells. Neurite outgrowth activity using rat pheochromocytoma (PC-12) cells was also determined. Twelve triterpenoids were isolated, of which several potently stimulated the expression of neurotrophic factors namely, *ngf* (sulphurenic acid, 15α-dehydroxytrametenolic acid, fomefficinic acid D and 16α-hydroxyeburicoic acid) and *bdnf* (sulphurenic acid and 15α- dehydroxytrametenolic acid), respectively. The triterpenes also potentiated *ngf*-induced neurite outgrowth in PC-12 cells. This is, to the best of our knowledge, the first report on the compound class of lanostanes in direct relation to *bdnf* and *ngf* enhancement. These compounds are widespread in medicinal mushrooms; hence, they appear promising as starting point for development of drugs and mycopharmaceuticals to combat neurodegenerative diseases. Interestingly, they do not show any pronounced cytotoxicity and may therefore be better suited for therapy than many other neurotrophic compounds that were previously reported.

## 1. Introduction

Neurodegenerative disease (NDD) is a term used to classify disorders that result in the progressive dysfunction of the nervous system such as Alzheimer’s disease and Parkinson’s disease [1,2]. These conditions lead to the loss and death of neural structure and function, which presents as deficits in different brain functions such as memory and cognition depending on the neurons affected by the disease. These conditions affect more than 50 million of the global population, with more than $600 billion required for the management of the condition [3]. Neurodegeneration may occur due to different biological mechanisms such as inflammation, oxidative stress, or mitochondrial dysfunction, among other factors. Current therapeutic interventions are concerned with relieving the symptoms of the disease and controlling the damage, although interventions that promote regeneration or offer neuron protection, and therapies that delay degeneration would be highly desirable [1,3].

Neurotrophins, such as Nerve Growth Factor (*ngf*) and Brain Derived Neurotrophic Factor (*bdnf*) are a group of proteins that help improve the survival of neurons, enhance their development and function in the central nervous system (CNS) [4,5]. While these proteins hold great promise in therapeutic applications, the challenge faced is that they have suboptimal pharmacological characteristics. For example, they are associated with poor serum stability, low bioavailability, and adverse effects, thus limited use in the management of NDD[6,7]. This indicates the need for different approaches, especially the use of neuroprotective strategies in managing NDD. The use of medicinal mushrooms is of great interest due to the potential therapeutic properties of metabolites and bioactive compounds [8].

Over the years, there has been intense exploration and investigation into the benefits of natural products and their bioactive components, especially those that have an NGF-potentiating activity with a low molecular weight or can mimic neurotrophic action and modulate signaling in the CNS [9,10]. These metabolites are preferred due to the ability to stimulate the production of neurotrophins, in addition to exerting antioxidative and anti-inflammatory properties, which provides neuroprotective action. Several studies showed that metabolites from *Hericium erinaceus* demonstrated neuritogenic properties in neuronal cells, which may be useful in the management of NDDs [6,11].

In our ongoing search for beneficial fungal metabolites, we have recently tested extracts and pure compounds from a wide range of other Basidiomycota and found some interesting hits. Like the aforementioned terpenoids from *Hericium*, these metabolites also induce *ngf* and *bdnf* mRNA levels on astrocytoma cells *in vitro* and effected neurite outgrowth activity using rat pheochromocytoma (PC-12) cells. The respective compounds were derived from fruitbodies of a German edible mushroom and the mycelial culture of an African polypore. The current paper is dedicated to describing the identity of the active principles. Since the two species showed similar metabolite profiles, we found it practical to combine the results of their evaluation in a single paper.

## 2. Results

### 2.1 Isolation of triterpenoids from L. sulphureus and Antrodia sp. (MUCL 56049)

A thorough analysis of the crude extracts from both *L. sulphureus* and MUCL 56049 *Antrodia* sp. by using high performance liquid chromatography–diode array detector high resolution mass spectrometry (HPLC–DAD–HRMS) suggested the presence of lanostane triterpenoids. Consequently, isolation and purification of the crude extracts by using reversed phase preparative RP-HPLC resulted to the elucidation of 12 known triterpenes (Figure 1). The structures were confirmed by comparison of their high resolution-electrospray ionization mass spectrometry (HR-ESIMS), their electrospray ionization mass spectrometry (ESIMS) and NMR spectroscopic data (Supplementary Information SI) with those reported in literature. The compounds from *L. sulphureus* were identified as laetiporin C or (3β,15α)-dihydroxy-24-hydroxymethyllanosta-8,23-dien-21-oic acid (**1**), laetiporin D or 15α-hydroxy-24-hydroxymethyl-3-oxolanosta-8,23-dien-21-oic acid (**2**) [12], trametenolic acid or 3β-hydroxylanosta-8,24-dien-21-oic acid (**3**) [13], eburicoic acid or 3β-hydroxy-24-methylene-lanost-8-en-21-oic acid (**4**) [14], 15α-hydroxytrametenolic acid or (3β,15α)-dihydroxylanosta-8,24-dien-21-oic acid (**5**) [15], sulphurenic acid or (3β,15α)-dihydroxy-24-methylene- lanost-8-en-21-oic acid (**6**) [16], fomefficinic acid D or 15α-hydroxy-24-methylene-3-oxolanosta-8-en21-oic acid (**7**) [17].

**Figure 1:**
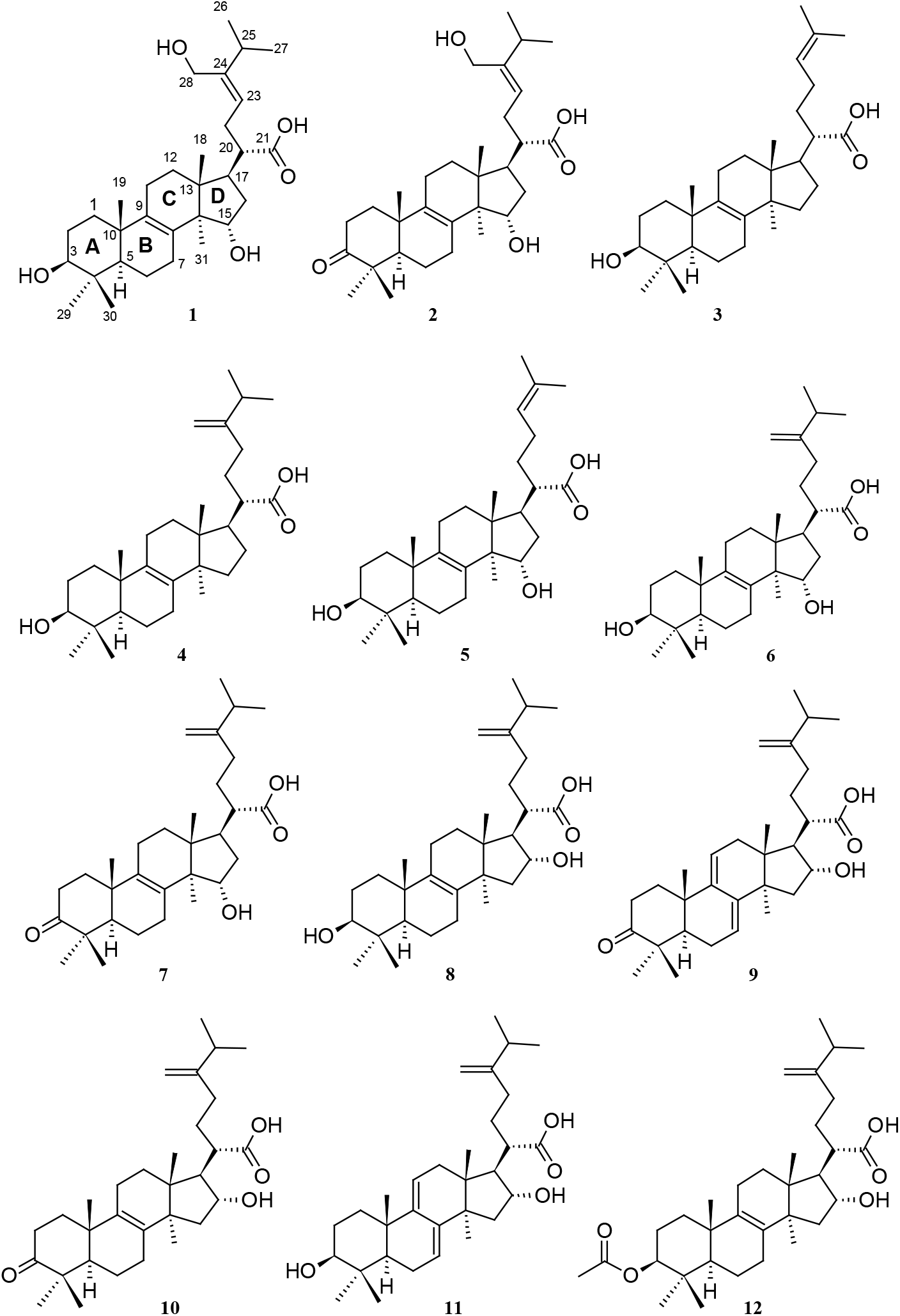
Isolated triterpenoids from *L. sulphureus* (**1**-**7**) and *Antrodia* sp. MUCL 56049 (**8**-**12)**

The metabolites from *Antrodia* sp. MUCL 56049 included tumulosic acid or (3β,16α)-dihydroxy-24-methylenelanost-8-en-21-oic acid (**8**) [18], polyporenic acid C or 16α-hydroxy-24-methylene-3-oxolanosta-7,9(11)-dien-21-oic acid (**9**) [19], 16α-hydroxyeburiconic acid or16α -hydroxy-24-methylene-3-oxolanost-8-en-21-oic acid (**10**) [19], dehydrotumulosic acid or (3β,16α)-dihydroxy-24-methylene- lanosta-7,9(11)-dien-21-oic acid (**11**) [20], and pachymic acid or 3β-acetyloxy- 16α-hydroxy-24-methylene-lanost-8-en-21-oic acid (**12**) [21].

### 2.2 Triterpenoids induce expression of ngf and bdnf in astrocytoma cells

Quantitative real-time reverse transcriptase polymerase chain reaction (RT-PCR) analysis revealed that treatments of the astrocytoma cell line 1321N1 cells with triterpenes (5 μg/mL) for 48h upregulate *ngf* and *bdnf* expression compared to DMSO treated cells (Figure 2). Strikingly, treatments with sulphurenic acid **6** (13-fold), 16α-hydroxyeburiconic acid **10** (5-fold), fomefficinic acid D **7** (4-fold) and 15α-dehydroxytrametenolic acid **5** (3-fold) significantly upregulate NGF mRNA expression levels while sulphurenic acid **6** (7-fold) and 15α- dehydroxytrametenolic acid **10** (4-fold) significantly upregulate *bdnf* expression. These results support the *ngf*-induced neurite outgrowth by PC-12 cells treated with triterpenoids isolated.

**Figure 2.**
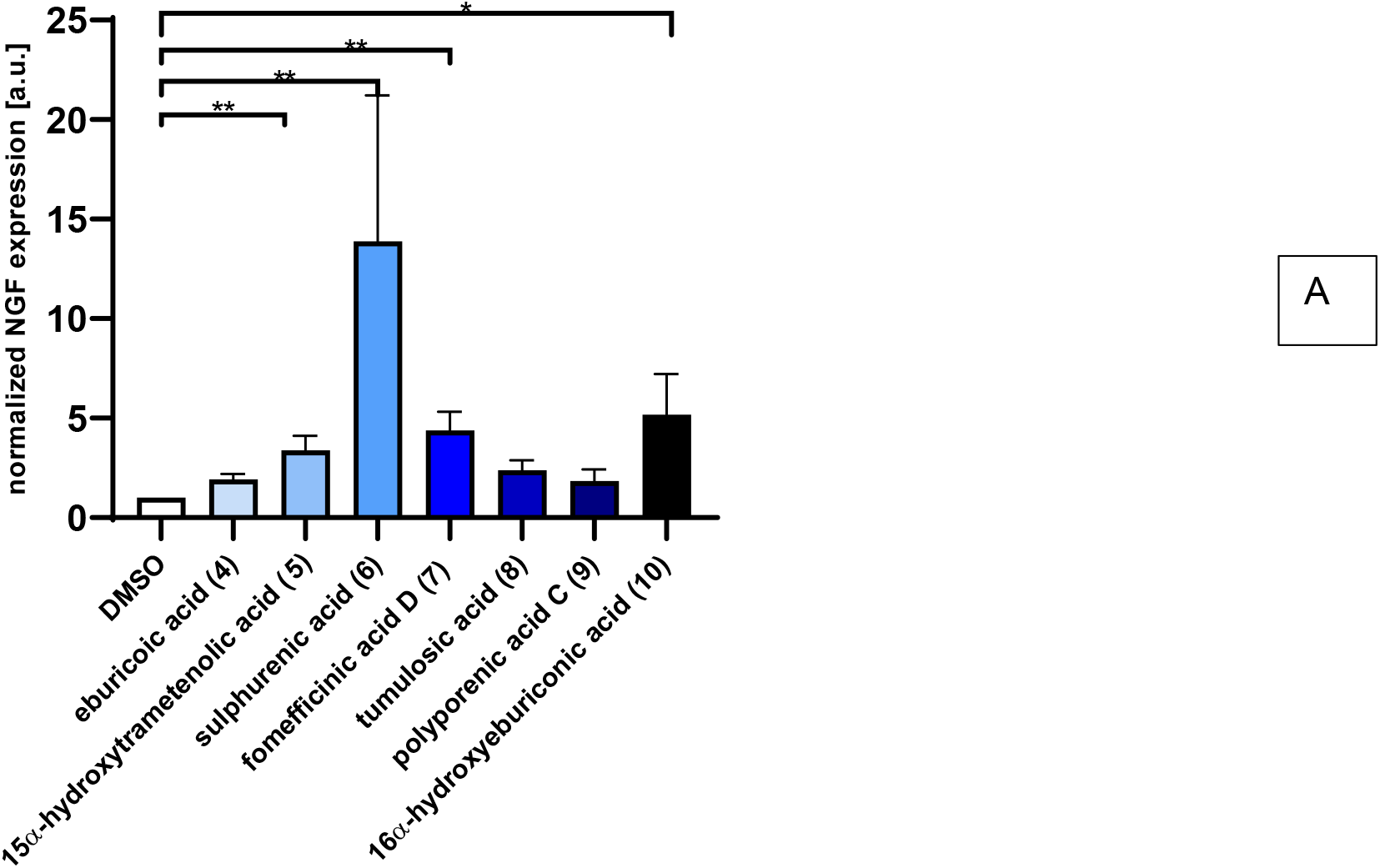
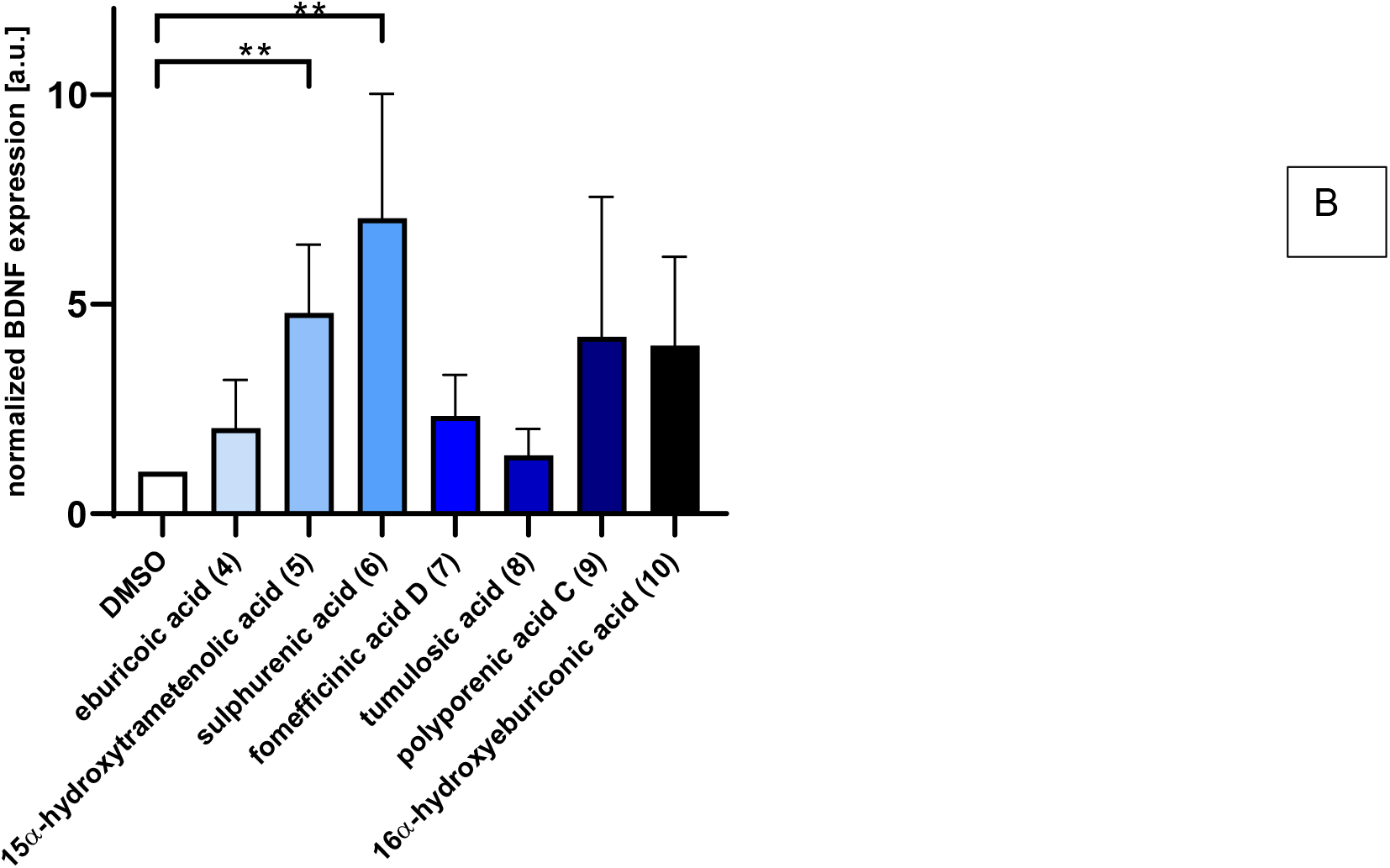
Triterpenoids induce both *ngf* and *bdnf* expression levels in 1321N1 astrocytoma cells. **A** and **B** Quantitative RT-PCR to compare the relative expression levels of *ngf* and *bdnf* in 1312N1 cells under triterpenes (5μg/mL) and 0.5% DMSO treatment for 48 h. Data shown as bar graphs are means ±S.E.M.; n = 3 to 5 independent experiments. *p<0.05, **p< 0.01, Mann-Whitney rank sum test.

Using cDNA synthesis and quantitative RT-PCR analysis, human astrocytoma 1321N1 cells treated with triterpenoids were found to upregulate the expression of genes for neurotrophins including nerve growth factor as well as brain derived neurotrophic factor – genes known to regulate neuroprotection [22]. According to [23], cyathane diterpenes isolated from the submerged cultures of *H. erinaceus* and *H. flagellum* were found to act on 1321N1 cells by increasing the transcription of either *bdnf* or *ngf*. Similarly, a study by [11] reported that erinacine C influenced the expression in these astrocytic cells on the transcriptional level, probably including also factors that regulate *ngf* expression. In the present study, we clearly demonstrated that the triterpenoids increases *ngf* and *bdnf* synthesis in 1321N1 cells. However, the amount of *ngf* secreted into the culture media after incubation with the compounds was not sufficient to promote the differentiation of PC-12 cells (preliminary work). This suggest that other neurotrophic factors play a role in the differentiation of PC-12 cells in addition to *ngf*.

### 2.3 Tritepenoids enhance ngf-induced neurite outgrowth in PC-12 cells

Measuring for neurite outgrowth of PC-12 cells is a well-established model [24] and the same methodology was also used in many previous studies on other fungal metabolites including our own. Hence, we tested lanostane triterpenes on the efficiency of neurite outgrowth and assessed the mean length of neurite outgrowth by live cell imaging after 72h (Figure 3). Neurites outgrowth were not observed when PC-12 cells were treated directly with the triterpenes. However, when PC-12 cells were treated with the triterpenes supplemented with 5ng/mL NGF, neurite outgrowth was observed (Figure 4). Treatment with sulphurenic acid (**6**), 15α-dehydroxytrametenolic acid (**5**), fomefficinic acid D (**7**) and eburicoic acid (**4**) caused a stronger differentiation of neurite bearing cells as compared to the control cells, which had 5 ng/mL of *ngf* alone (Figure 5) over time. Investigations whether the above mentioned triterpenes potentiate *ngf*-induced neurite outgrowth by stimulating *ngf* synthesis or as substitutes for *ngf* (*ngf*-mimicking activity) are ongoing.

**Figure 3.**
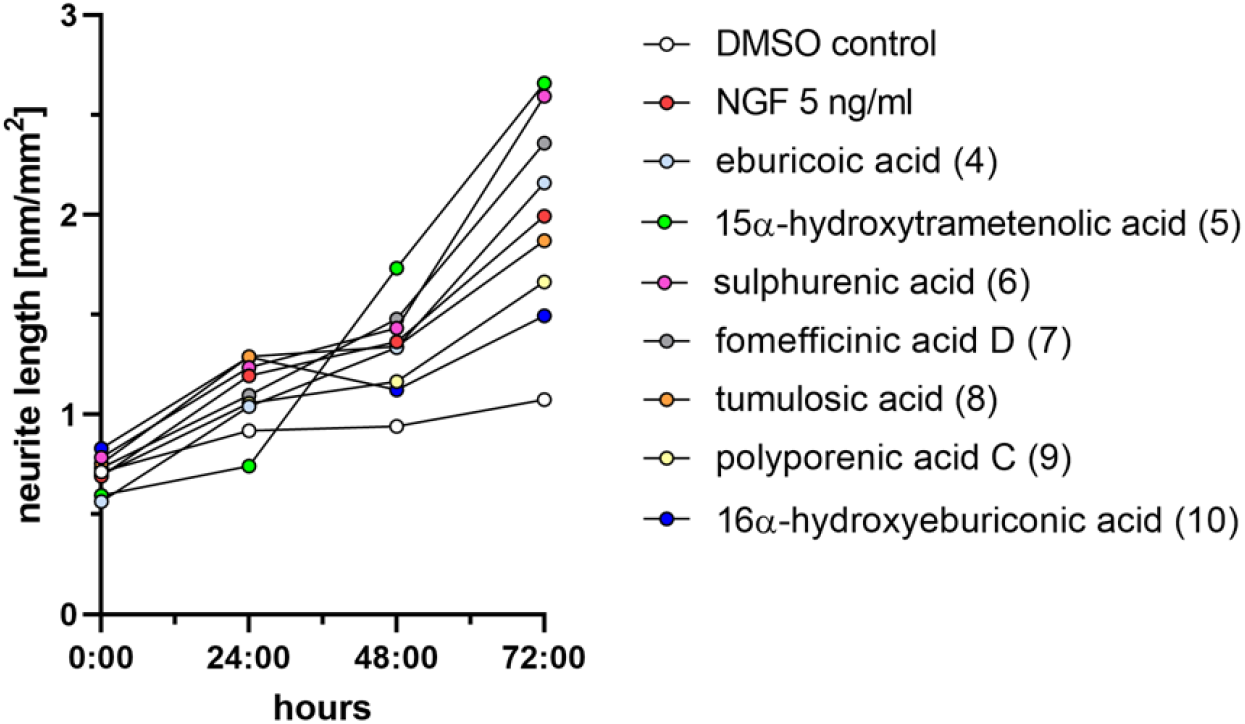
Graph showing the mean neurite length of PC-12 cells treated with different triterpenes supplemented with 5ng/mL NGF over a time period of 72h. Negative control is PC-12 cell treated with 0.5% DMSO. Positive control is 5ng/mL NGF.

**Figure 4.**
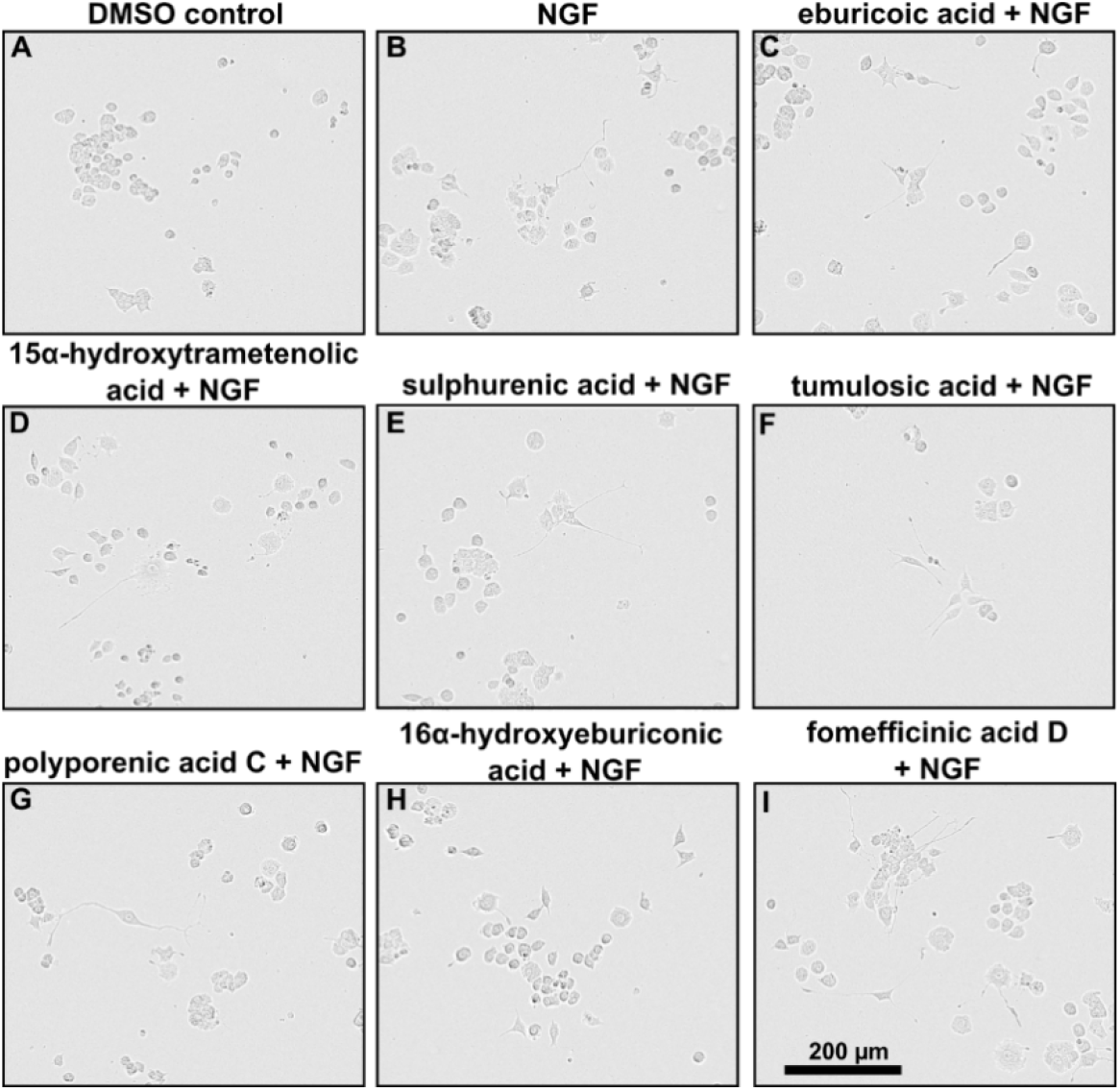
(A-I) PC-12 cells treated with 5 ng/mL NGF and either DMSO or different triterpenes (5 μg/mL) were recorded for 3 days. Phase contrast images of cells after 3 days of treatment.

**Figure 5.**
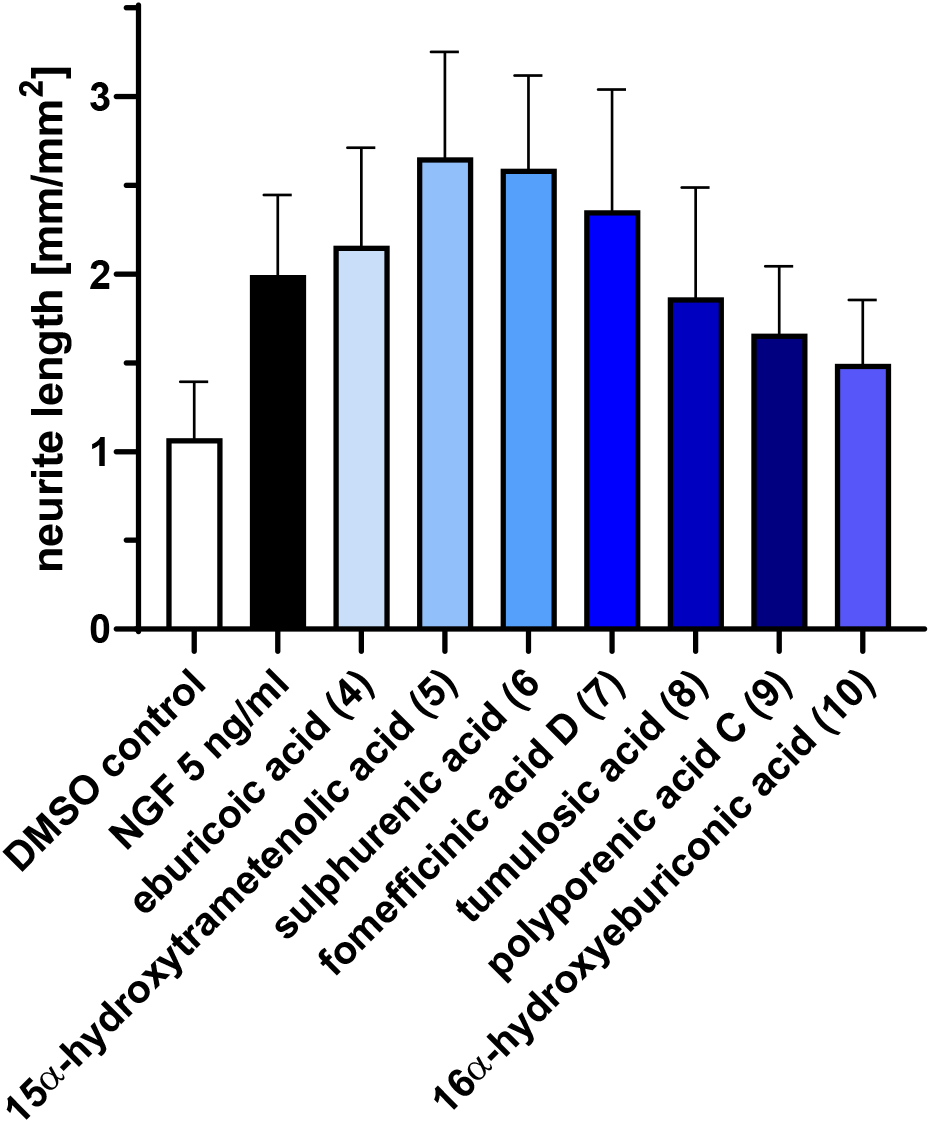
Graph showing the mean neurite length of PC-12 cells treated with different triterpenes supplemented with 5ng/mL NGF after 72 h ± S.E.M. Data originate from three independent experiments.

## 3. Discussion

The results of our current study have highly interesting implications for the potential utility of lanostane triterpenoids, which are very commonly found in wood-inhabiting Basidiomycota, including many species of medicinal mushrooms that have been used for millennia in traditional Asian medicine. We will discuss some aspects that relate to the current context.

### 3.1 Previous reports of neurotrophic activities of extracts derived from other Basidiomycota and selected reports on neurotropic effects of plant metabolites

In our study, several triterpenes supplemented with *ngf* exhibited enhanced neurite outgrowth of PC-12 cells compared to *ngf* alone. This observation is in agreement with other studies. Report by [25] show a remarkably increased neurite outgrowth on PC-12 cells treated with hericenones C, D and E isolated from fruiting bodies of *H. erinaceus* in the presence of 5 ng/mL *ngf*, and with further references on *Hericium* spp. cited earlier in this paper.

Similarly, [26] observed no cell differentiation or neurite outgrowth in PC-12 cells treated with littorachalcone, isolated from *Verbena littoralis* in the absence of *ngf*, but markedly enhanced induced neurite outgrowth activity in the presence of *ngf* (2 ng/mL). A study by [27] reported that a combination of *ngf* and sclerotium extract of the Malaysian medicinal mushroom *Lignosus rhinoceritis* had additive effects and enhanced neurite outgrowth. On the other hand, reports on neurite outgrowth on PC-12 cells treated with both, crude extracts and pure compounds without the presence of *ngf* are documented [5,28,29]. However, none of the aforementioned papers has been explicitly dealing with a lanostane triterpenoid that was identified as active principle. These studies either dealt with non-defined crude extracts that were not even characterized analytically, or different metabolites were shown to be responsible for the observed biological effects. The study by [5] reported another plant-derived triterpenoid of a different type (cucurbitane), lindersin B, to be active as NGF enhancer by activation of the tyrosine kinase A/phosphatidylinositol 3 kinase/extracellular signal-regulated kinase signaling pathways. [30] also demonstrated the neurite outgrowth-promoting activity of hopane-type triterpenoid 2-acetoxydiplopterol isolated from the leaves of *Illicium merrillianum* using PC-12 cells. The neurotrophic activities of several other plant-derived tripterpenes of different types including oleanane-type (oleanolic acid), and ursane-type (ursolic acid) have also been reported. The latter compounds showed a markedly enhancing activity of *ngf*-mediated neurite in PC-12D cells [31]. Oleanolic acid was also proven to ameliorate scopolamine-induced memory impairment by modulating the *bdnf-* ERK1/2-CREB pathway through TrkB activation in mice [32].

### 3.2 Known biological activities of compounds 1 – 12 and related triterpenes

Sulphurenic acid (**6**) was one of the compounds that showed the most significant effect in our studies and affected both, *bdnf* and *ngf* enhancement. The compound was first identified by [16] from basidiomata of *“Polyporus”* (i.e., *Laetiporus sulphureus*) and has since then been discovered several times from various species of *Laetiporus* and other members of the *Polyporales* [33,34], including *“Antrodia camphorata”* (i.e. *Taiwanofungus camphoratus*) [35,36]. In their study of the constituents of the latter fungus, [37] noted only very weak cytotoxic effects, which is in agreement with our own findings, reported by [12,14]. As the compound does not apparently affect mammalian cells so strongly but has the strongest neurotrophic effects, it may be one of the most promising candidates for further evaluation. Interestingly, it has also been discovered in *Fomitopsis officinalis* [38], also known under the synonyms *Fomes* or *Laricifomes* which is one of the most well-known medicinal mushrooms that have been used in Asia as well as in Europe for a very long time [39].

15α-dehydroxytrametenolic acid (**5**), the second compound that strongly enhanced *bdnf* and *ngf* production, as well as fomefficinic acid D (**7**), and 16α-hydroxyeburicoic acid (**10**), which only had a significant effect on *ngf* production, are also known from this group of fungi and have often been encountered concurrently with the aforementioned compounds [17,33,37].

Pachymic acid (**12)** has first been reported by [21] from basidiomata of the medicinal mushrooom *“Poria cocos”* (currently valid name: *Wolfiporia cocos*), which also belongs to the Fomitopsidaceae. No bioactivities were initially reported, but the compound has later on found to be active in various studies, relating to its anticancer and anti-inflammatory properties [40,41]. It was not among the strongest NGF enhancers that we found in the current study. There are also additional derivatives of related metabolites, including some that have even been isolated and characterized from *Laetiporus* species. For example, an acetyl derivative of eburicoic acid (**4**) has also been reported to have anti-inflammatory effects on murine macrophages [42].

[43,44] reported that triterpenoids isolated from *“Ganoderma lucidum”* showed neurotropic properties by enhancing the survival ad promoting the survival of neurons. These authors, however, did not show these effects in a direct manner, but tested their compounds for their ability to initiate TrkA- or TrkB-related signal transduction in NIH-3T3/TrkA and NIH-3T3/TrkB cells were examined using an MTT assay. The two derivatives, 4,4,14α-trimethyl-5α-chol-7,9(11)-dien-3- oxo-24-oic acid (which was then reported to be a novel natural derivative) and methyl ganoderic acid B, and six other congeners did not show such effects or were not examined. In addition, triterpenes isolated from *“G. lucidum“* have also been reported to attenuate the production of pro-inflammatory cytokines in macrophage cells [45]. The fungus that these authors have reported on, however, is probably not identical with the European species *G. lucidum*, which does not occur in China where these studies were undertaken [46]. It may correspond with *G. lingzhi* (i.e. the fungus that was confused with *G. lucidum* and has been used in Traditional Asian medicine for centuries) or another native Chinese species (cf.) [47]. Notably, many of the triterpenes from *Ganoderma* species are rather cytotoxic (and preparations made from *Ganoderma* are even in use as anticancer agents). In contrast, the compounds that were identified as neurotrophins in the current study have only weak cytotoxic effects on mammalian cells [12,14] and are also lacking certain structural features like the carboxyl group at C-24 of the triterpenoid skeleton. Even though the *Ganoderma* triterpenoids were apparently not yet tested for NGF-enhancing activities in our bioassays, it seems possible that the ones from *Antrodia* and *Laetiporus* exert their neurotrophic effects through a different mode-of-action. Interestingly, these triterpenoids are also more selective neurotrophic accents than e.g. the diterpenoids of the erinacine type from *Hericium*, some of which show cytotoxicity in the low μM range (cf.) [23]. On the other hand, the triterpenoids exhibit their neurotrophic effects at much lower concentrations than the corallocins from *Hericium coralloides* and related meroterpenoids that are also present in *H. erinaceus* (cf.) [48].

### 3.3 Previous use of Laetiporus and Antrodia species and their relatives in traditional medicine and for other applications

The genus *Laetiporus* belongs to the Fomitopsidaceae (Polyporales) and its species form conspicuous basidiomes with bright yellowish colors on wood. They are considered to be forest pathogens and cause brown rot, but some are edible and medicinal mushrooms [49]. The genus is geographically distributed world-wide and its species can be found in cold temperate to tropical zones. The type species, *Laetiporus sulphureus*, goes back to the French mycologist Buillard, who described it as *Boletus sulphure*us in 1789, and is very frequently encountered in Europe and other areas of the Northern hemisphere. It is known as “Chicken of the woods”, or “sulphur shelf” in English speaking countries and as “Schwefelporling” in Germany. Out of the 15 presently accepted species, eleven belong to the *L. sulphureus* complex, as was recently established by phylogenetic analyses [50,51].

The young fruitbodies are of soft consistence and are even regarded as a culinary delicacy. In China and Europe the fruitbodies have been used traditionally to strengthen the human body and as remedy for various diseases (see summary by [52]). However, according to our literature search, no study has yet been undertaken to link the reported health benefits directly and conclusively to the triterpenoids and other secondary metabolites of *L. sulphureus*.

Like *Laetiporus*, the genus *Antrodia* also belongs to the Fomitopsidaceae and is typified by *A, serpens*, a fungus that was first described by [53] as *Polyporus serpens*. The genus has presently more than 50 accepted species [54,55], which are also known to destroy wood by causing brown rot.

The taxonomy of MUCL 56049 is also still unclear. The genus was determined from morphological traits and because the closest matches retrieved from a BLAST search in GenBank based on LSU and ITS sequences were derived from a specimen of *“Antrodia* aff. *albidoides”* sensu Ryvarden from Zimbabwe, which has likewise not been formally described. The ITS sequence (GenBank acc no KC543176.1) has been generated and uploaded by [55] and also shows high homologies to other species in the family Fomitopsidaceae, to which the genera *Laetiporus* and *Fomitopsis* belong. Many of the tropical species of *Antrodia* and allied genera, especially from Africa remain widely uncharacterized by modern polyphasic taxonomic methodology, and a recent monograph is not available, which is why we have thus far been unable to classify the specimen MUCL 56049 to species level.

Regarding the practical use of *Antrodia* species as food or medicinal mushroom, there is not much information in the literature. The fruitbodies are inedible, due to their tough consistency, and the reports on medicinal usages of *“Antrodia”* species can mostly be related to *Taiwanofungus camphoratus*, an important Asian medicinal mushroom related to the species of *Antrodia*, which has in the past been referred to under the synonyms *A. camphorata* or *A. cinnamomea* [17]. The economic value of *T. camphoratus* is mainly due to its medicinal use and it belongs to the most valued medicinal fungus in Asian countries. This species has already been used as a natural therapeutic ingredient in Traditional Chinese Medicine (TCM) for its antioxidative, anti-tumor, anti-cancer, anti-hepatitis, vasorelaxation, anti-inflammatory, cytotoxic, and neuroprotective activities [33,54,56].

Because of its use in traditional medicine, the secondary metabolites of *T. camphoratus* have been studied intensively, and over 70 compounds have been elucidated from this fungus, 39 of which are triterpenoids [54,56]. These triterpenes are of the ergostane and lanostane types (tumulosic acid (**8**), polyporenic acid C (**9**), 16α-hydroxyeburicoic acid (**10**), dehydrotumulosic acid (**11**) and pachymic acid (**12**) that were also found in this study from the African *Antrodia* sp. Neuroprotective effects have been claimed in two patent applications [57,58] in which some of the compounds we studied were mentioned, even though no direct evidence has been provided by these authors, and no NGF or BDBF enhancing activities were noted in these patent applications. On the other hand, the current study clearly shows that it is worthwhile to further study these fungi, including their secondary metabolism, as this may reveal interesting chemotaxonomic relationships as well as hitherto unprecedented biological activities of their constituents.

## 4. Materials and Methods

### 4.1 General Information

HPLC-DAD/MS measurements were performed using an amaZon speed ETD (electron transfer dissociation) ion trap mass spectrometer (Bruker Daltonics, Bremen, Germany) and measured in positive and negative ion modes simultaneously. The HPLC system comprised the following elements; Column C18 Acquity UPLC BEH (Waters); mobile phase: solvent A (H2O)/solvent B: acetonitrile (ACN), supplemented with 0.1% formic acid; gradient conditions: 5% B for 0.5 min, increasing to 100% B 20, maintaining isocratic conditions at 100% B for 10 min, flow rate 0.6 mL/min, UV/Vis detection 200–600 nm).

HRESIMS (high-resolution electrospray ionization mass spectrometry) data were recorded on a mass spectrometer (Bruker Daltonics,) coupled to an Agilent 1260 series HPLC-UV system and equipped with C18 Acquity UPLC BEH (ultraperformance liquid chromatography) (ethylene bridged hybrid) (Waters) column; DAD-UV detection at 200–600 nm; solvent A (H2O) and solvent B (ACN) supplemented with 0.1% formic acid as a modifier; flowrate 0.6 mL/min, 40 °C, gradient elution system with the initial condition 5% B for 0.5 min, increasing to 100% B in 19.5 min and holding at 100% B for 5 min. To determine the molecular formula, Compass DataAnalysis 4.4 SR1 was used using the Smart Formula algorithm (Bruker Daltonics). NMR spectra were recorded on a Bruker 700 MHz Avance III spectrometer equipped with a 5 mm TCI cryoprobe (1H: 700 MHz, 13C: 175 MHz), locked to the respective deuterium signal of the solvent.

### 4.2 Fungal material

The species that were studied on the production of neurotrophins were selected out of our ongoing research program for discovery of antibiotics and other beneficial metabolites from European and tropical Basidiomycota. In this scenario, samples (extracts and pure compounds) that are devoid of cytotoxic and prominent antimicrobial effects are being investigated for other potential applications. The current study deals with a fruitbody extract of specimens collected in Germany and an extract derived from a mycelial culture of an African species.

*Laetiporus sulphureus* (Figure 6A) was collected in Braunschweig-Riddagshausen nature reserve on 27 May, 2020. Hosts plants were *Salix* spec., *Fagus sylvatica*, and *Robinia pseudoacacia*. The specimens were identified by Harry Andersson based on morphological data and a voucher specimen was deposited in the fungarium of the HZI, Braunschweig. Further details, including the isolation of the terpenoids that were tested in this study, have recently been described by [12].

The fruitbodies of *Antrodia* sp. (Figure 6B) were collected by C. Decock and J. C. Matasyoh from Mount Elgon National Reserve (1°7’6” N, 34°31’30’ E) in Kenya in April 2016. A dried specimen 00and a corresponding culture were deposited at Mycothèque de l’Université catholique de Louvain (MUCL, Louvain-la-Neuve, Belgium) under the accession number MUCL56049. The genus was identified by morphological studies and comparison of the internal transcribed spacer (ITS)-nrDNA sequences with others from Basidiomycota deposited in GenBank. The sequence is also available in the Supplementary Information.

### 4.3 Extraction and isolation of the secondary metabolites

The extraction and purification of compounds from *L. sulphureus* was described in detail in our recent paper [12] and led to isolation of seven metabolites (**1**-**7**). The same protocols were used for isolation of the compounds from a batch culture of *Antrodia* sp. MUCL 56049 that had been grown in twenty 500 mL shake flasks containing each 200 mL of yeast-malt (YM 6.3) medium (10 g/L malt extract, 4 g/L yeast extract, 4 g/L D-glucose and pH = 6.3) for 24 days. The mycelia were harvested three days after glucose depletion by filtration and the culture filtrate was set aside as it did not contain any target metabolites.

Extraction of the mycelia of *Antrodia* sp. from the aforementioned four litres of culture media with acetone (200mL) led to a total of 442 mg crude product. The crude extract (cream solid) was dissolved in methanol (MeOH) and filtered using cotton in a pasteur pipette. The crude extract was fractionated using preparative reverse phase liquid chromatography (PLC 2020, Gilson, Middleton, USA). VP Nucleodur 100-5C 18 ec column (250 ×40 mm, 7 μm: Macherey-Nagel) used as stationary phase. Deionized water (Milli-Q, Millipore, Schwalbach, Germany) (solvent A) and acetonitrile (solvent B) with 0.05% TFA was used as eluent with flow rate of 40 mL/min. Three runs were performed on the preparative RP-HPLC with the elution gradient of 50 - 100% solvent B in 60 min and thereafter isocratic condition at 100% solvent B for 5 min. UV/Vis detection was carried out at 210nm and 300 nm. Eleven fractions were collected according to the selected peaks. Similar fractions from each round of separation were combined consequently leading to isolation of compounds **8** (4 mg), **9** (1.25 mg), **10** (5.2 mg), **11** (2.2 mg), and **12** (3.5 mg). Their structures were elucidated by using their HR-ESIMS data and their 1D and 2D NMR spectroscopic data, which were then compared with those already reported in the literature. The spectral data for compounds **1**-**7** are deposited in the Supplementary Information with the preceding studies by [12,14] and the data for compounds **8**-**12** are given in the Supplementary Information of the current paper.

### 4.4 Cell Culture

Astrocytoma (1321N1) cells (obtained from Sigma-Aldrich, acc. No. 86030402) were cultured in Gibco DMEM medium (Fisher Scientific, Inc., Waltham, MA, USA) containing 10% heat-inactivated FBS (Capricorn) [11]. Rat pheochromocytoma cells (PC-12, adherent variant) purchased from European Collection of Authenticated Cell Cultures (ECACC) general collection were grown in Gibco™ RPMI-1640 (Fisher Scientific, Hampton, NH, USA) medium containing 10% horse serum (Capricorn™ Scientific GmbH, Ebsdorfergrund, Germany) and 5% heat-inactivated fetal bovine serum-FBS (Capricorn). The media were supplemented with penicillin (0.15 mM), streptomycin (86 μM), and glutamine (2 mM) [23]. The cells were incubated at 37°C in a humidified environment of 7.5% CO2 and were routinely passaged every 3-4 days. Collagen type IV (Sigma C5533) was coated on 96 well plates and left for 6 hours or more before using the plates whenever seeding PC-12 cells.

### 4.5 cDNA synthesis and real-time quantitative RT-PCR

Since PC-12 cells do not produce *ngf* by their own, confirming directly the induction of neurotrophin expression in 1321N1 cells was tested. For real-time quantitative reverse transcriptase PCR the total RNA was extracted from triterpene treated 1321N1 cells (2×10^5^ cells). For treatment, culture media was replaced by serum-reduced medium (Gibco RPMI with 1% FBS (Capricorn)) and cells were incubated for 24h. Then, media was replaced with media containing the triterpenes dissolved in 0.5% DMSO. As control, serum-reduced medium supplemented with 0.5% DMSO was used. Cells were incubated for 48 h [11,23]. Total RNA was extracted using the NucleoSpin^R^ RNA Plus kit (Macherey-Nagel GmbH& Co KG, Germany) followed by further purification (NucleoSpin^R^ RNA Clean-up kit) according to the manufacturer’s protocol. To determine the concentration of purified RNA the corresponding samples were measured using a DS-11+ spectrophotometer Nanodrop (DeNovix Inc. Wilmington, Delaware USA). First-strand cDNA synthesis and subsequent real-time PCR was performed using SensiFast ™ SYBR No-Rox One-Step Kit (Cat.No. BIO- 72005 (Bioline)). The following PCR primers were used for amplifying specific cDNA fragments: *gapdh* (sense 5’-ACCACAGTCCATGCCATCAC-3’; antisense: 5‘- TCCACCACCCTGTTGCTGTA-3’ 451 bp), *ngf* (sense: 5‘- CCAAGGGAGCAGTTTCTATCCTGG-3’; antisense 5‘- GGCAGTTGTCAAGGGAATGCTGAAGTT-3’ 189 bp) and *bdnf* (sense: 5’- TAACGGCGGCAGACAAAAAGA-3’; antisense: 5’-GAAGTATTGCTTCAGTTGGCCT-3‘; 101 bp) [11]. The PCR reactions were performed in a 10 μL volume containing cDNA template (2 μL), SensiFast ™ SYBR No-Rox One-step Mix (5 μL), primers (400 nM; 0.4 μL), Reverse transcriptase (0.1 μL), RiboSafe RNase Inhibitor (0.2 μL) and Rnase free water (1.9 μL). The amplified cDNAs were analyzed and quantified using the Qiagen (Corbett) Rotor–Gene ™ 3000 and LightCycler^R^ 96 (Roche Diagnostics International Ltd, version 1.1.0.1320) real time PCR instruments. Amounts of *gapdh* amplicons value were used as reference and set as 1.

### 4.6 Neurite outgrowth assay

Preliminary studies had shown that conditioned media from 1321N1 cells treated with the isolated triterpenes did not display PC-12 cell differentiation activity triggering neurite outgrowth. The neurotrophic activity assay was conducted as described by Phan et al. (2015). PC-12 cells were seeded at a density of 1×10^3^ cells per well in growth medium in 96- well culture plates and incubated overnight. Triterpenoids isolated from *L. sulphureus* and *Antrodia* sp. were added directly to the cells and further incubated at 37°C with 5% CO2 for 3 days. Cells treated with nerve growth factor (50ng/mL) were used as a positive control. After 3 days, the cells were examined using an IncuCyte S3 live-cell analysis system (Sartorius, Göttingen, Germany). Six random fields were examined in each well. The number of neurite outgrowth (axon-like processes), defined as extensions longer than twice the cell body diameter, was recorded. Neurite length was measured using the IncuCyte NeuroTrack Software Module (2019B Rev2 GUI) software. Three independent experiments were conducted for each compound. Neurite outgrowth activity in the presence of both *ngf* and triterpenoids was also then analyzed. A concentration of *ngf* (5 ng/mL) was added in addition to the triterpenoids on the PC-12 cells, and the neurite outgrowth recorded. Three independent experiments were performed in this assay.

### 4.7 Statistical analysis

Data are displayed as the mean ± SEM and analyzed by Mann-Whitney rank sum test using the software Prism V8 (Graphpad Software Inc., San Diego, CA, USA).

## 5. Conclusions

What remains to be done is to establish a clear-cut structure-activity relationship between the substitution patterns of the lanostane triterpenoids and the corresponding biological effects. Since it was not possible to establish this based on the relatively low number of natural products available as now, it may be feasible to isolate large quantities of one of the most potent derivatives and subject this metabolite to semi-synthesis. On the other hand it could be feasible to use the mixtures of triterpenoids that can be enriched by the development of special extraction techniques, as a standardized product that can be used as a “mycopharmaceutical” or, in case of the edible *Laetiporus*, as a nutraceutical that can be used as prophylactic agent for prevention of neuropathic disorders. In any case, the biochemical mode of action of these compounds also needs further study because so far it is not evident how the neurotrophic effects observed actually came about. However, the amounts of compounds that became available during the course of this work were too small to conduct follow up experiments as described above and need to follow in the near future after further scale up of production and isolation of the terpenoids. For this purpose, it may even be feasible to study related fungi in order to get better yields, or to optimize the fermentation process in view of a transfer to larger scale bioreactors.

## Supporting information

Total supplemental info

## Supplementary Materials

The following supporting information can be downloaded at: www.mdpi.com/xxx/s1, Figure S1: ESIMS data for tumulosic acid (8); Figure S2: HR-ESIMS data for tumulosic acid (8); Figure S3: 1H NMR spectrum (DMSO-*d_6_*, 700 MHz) of tumulosic acid (8); Figure S4: COSY spectrum (DMSO- *d_6_*, 700 MHz) of tumulosic acid (8); Figure S5: HSQC spectrum (DMSO- *d_6_*, 700 MHz) of tumulosic acid (8); Figure S6: HMBC spectrum (DMSO- *d_6_*, 700 MHz) of tumulosic acid (8); Figure S7: ESIMS data for polyporenic acid C (9); Figure S8: HR-ESIMS data for polyporenic acid C (9); Figure S9: 1H NMR spectrum (DMSO- *d_6_*, 700 MHz) of polyporenic acid C (9); Figure S10: COSY spectrum (DMSO- *d_6_*, 700 MHz) of polyporenic acid C (9); Figure S11: HSQC spectrum (DMSO- *d_6_*, 700 MHz) of polyporenic acid C (9); Figure S12: HMBC spectrum (DMSO- *d_6_*, 700 MHz) of polyporenic acid C (9); Figure S13: ESIMS data for 16α-hydroxyeburiconic acid (10); Figure S14: HR-ESIMS data for 16α-hydroxyeburiconic acid (10); Figure S15: 1H NMR spectrum (DMSO- *d_6_*, 700 MHz) of 16α-hydroxyeburiconic acid (10); Figure S16: COSY spectrum (DMSO- *d_6_*, 700 MHz) of 16α-hydroxyeburiconic acid (10); Figure S17: HSQC spectrum (DMSO- *d_6_*, 700 MHz) of 16α-hydroxyeburiconic acid (10); Figure S18: HMBC spectrum (DMSO- *d_6_*, 700 MHz) of 16α-hydroxyeburiconic acid (10); Figure S19: ROESY spectrum (DMSO- *d_6_*, 700 MHz) of 16α-hydroxyeburiconic acid (10); Figure S20: ESIMS data for dehydrotumulosic acid (11); Figure S21: HR-ESIMS data for dehydrotumulosic acid (11); Figure S22: 1H NMR spectrum (DMSO- *d_6_*, 700 MHz) of dehydrotumulosic acid (11); Figure S23: COSY spectrum (DMSO- *d_6_*, 700 MHz) of dehydrotumulosic acid (11); Figure S24: HSQC spectrum (DMSO- *d_6_*, 700 MHz) of dehydrotumulosic acid (11); Figure S25: HMBC spectrum (DMSO- *d_6_*, 700 MHz) of dehydrotumulosic acid (11); Figure S26: ESIMS data for pachymic acid (12); Figure S27: HR-ESIMS data for pachymic acid (12); Figure 28: 1H NMR spectrum (DMSO- *d_6_*, 700 MHz) of pachymic acid (12); Figure S29: COSY spectrum (DMSO- *d_6_*, 700 MHz) of pachymic acid (12); Figure S30: HSQC spectrum (DMSO- *d_6_*, 700 MHz) of pachymic acid (12); Figure S31: HMBC spectrum (DMSO- *d_6_*, 700 MHz) of pachymic acid (12); Figure S32: ROESY spectrum (DMSO- *d_6_*, 700 MHz) of pachymic acid (12); ITS and LSU sequences of Antrodia sp.

## Author Contributions

Conceptualization: Marc Stadler, Reinhard Köster; Methodology: Marc Stadler, Reinhard Köster, Theresia E.B. Stradal; Formal analysis and investigation: Kathrin Wittstein, Monique Rascher, Clara Chepkirui, Khadija Hassan, Marco Kirchenwitz, Blondelle Matio Kemkuignou; Writing - original draft preparation: all authors; Writing - review and editing: All authors. Funding acquisition: Marc Stadler, Reinhard Köster, Theresia E.B. Stradal, Josphat C. Matasyoh, Cony Decock. All authors read and approved the final manuscript.

## Data Availability Statement

All data generated or analyzed during this study are included in this published article and its supplementary information files.

## Acknowledgments

We are grateful to Wera Collisi, Christel Kakoschke Silke Reinecke and Silvia Pettrin for their technical support and to Harry Anderson for collection of *Laetiporus* specimen.

