## Supplementary material for "Neurotrophic and immunomodulatory lanostane triterpenoids from wood-inhabiting Basidiomycota": Total supplemental info

Table of Contents

**Figure S1**: ESIMS data for tumulosic acid (**8**). 3

**Figure S2**: HR-ESIMS data for tumulosic acid (**8**). 4

**Figure S3**: $^{1}$H NMR spectrum (DMSO-$d_6$, 700 MHz) of tumulosic acid (**8**). 5

**Figure S4**: COSY spectrum (DMSO-d6, 700 MHz) of tumulosic acid (**8**). 6

**Figure S5**: HSQC spectrum (DMSO-d6, 700 MHz) of tumulosic acid (**8**). 7

**Figure S6**: HMBC spectrum (DMSO-$d_6$, 700 MHz) of tumulosic acid (**8**). 8

**Figure S7**: ESIMS data for polyporenic acid C (**9**). 9

**Figure S8**: HR-ESIMS data for polyporenic acid C (**9**). 10

**Figure S9**: $^{1}$H NMR spectrum (DMSO-$d_6$, 700 MHz) of polyporenic acid C (**9**). 11

**Figure S10**: COSY spectrum (DMSO-$d_6$, 700 MHz) of polyporenic acid C (**9**). 12

**Figure S11**: HSQC spectrum (DMSO-$d_6$, 700 MHz) of polyporenic acid C (**9**). 13

**Figure S12**: HMBC spectrum (DMSO-$d_6$, 700 MHz) of polyporenic acid C (**9**). 14

**Figure S13**: ESIMS data for 16α-hydroxyeburiconic acid (**10**). 15

**Figure S14**: HR-ESIMS data for 16α-hydroxyeburiconic acid (**10**). 16

**Figure S15**: $^{1}$H NMR spectrum (DMSO-$d_6$, 700 MHz) of 16α-hydroxyeburiconic acid (**10**). 17

**Figure S16**: COSY spectrum (DMSO-d6, 700 MHz) of 16α-hydroxyeburiconic acid (**10**). 18

**Figure S17**: HSQC spectrum (DMSO-$d_6$, 700 MHz) of 16α-hydroxyeburiconic acid (**10**). 19

**Figure S18**: HMBC spectrum (DMSO-$d_6$, 700 MHz) of 16α-hydroxyeburiconic acid (**10**). 20

**Figure S19**: ROESY spectrum (DMSO-$d_6$, 700 MHz) of 16α-hydroxyeburiconic acid (**10**). 21

**Figure S20**: ESIMS data for dehydrotumulosic acid (**11**) 22

**Figure S21**: HR-ESIMS data for dehydrotumulosic acid (**11**). 23

**Figure S22**: $^{1}$H NMR spectrum (DMSO-$d_6$, 700 MHz) of dehydrotumulosic acid (**11**). 24

**Figure S23**: COSY spectrum (DMSO-$d_6$, 700 MHz) of dehydrotumulosic acid (**11**). 25

**Figure S24**: HSQC spectrum (DMSO-$d_6$, 700 MHz) of dehydrotumulosic acid (**11**). 26

**Figure S25**: HMBC spectrum (DMSO-$d_6$, 700 MHz) of dehydrotumulosic acid (**11**). 27

**Figure S26**: ESIMS data for pachymic acid (**12**) 28

**Figure S27**: HR-ESIMS data for pachymic acid (**12**) 29

**Figure 28**: 1H NMR spectrum (DMSO-d6, 700 MHz) of pachymic acid (**12**). 30

**Figure S29**: COSY spectrum (DMSO-$d_6$, 700 MHz) of pachymic acid (**12**). 31

**Figure S30**: HSQC spectrum (DMSO-$d_6$, 700 MHz) of pachymic acid (**12**). 32

**Figure S31**: HMBC spectrum (DMSO-$d_6$, 700 MHz) of pachymic acid (**12**). 33

**Figure S32**: ROESY spectrum (DMSO-$d_6$, 700 MHz) of pachymic acid (**12**). 34

**ITS and LSU sequences of *Antrodia* sp.** 35

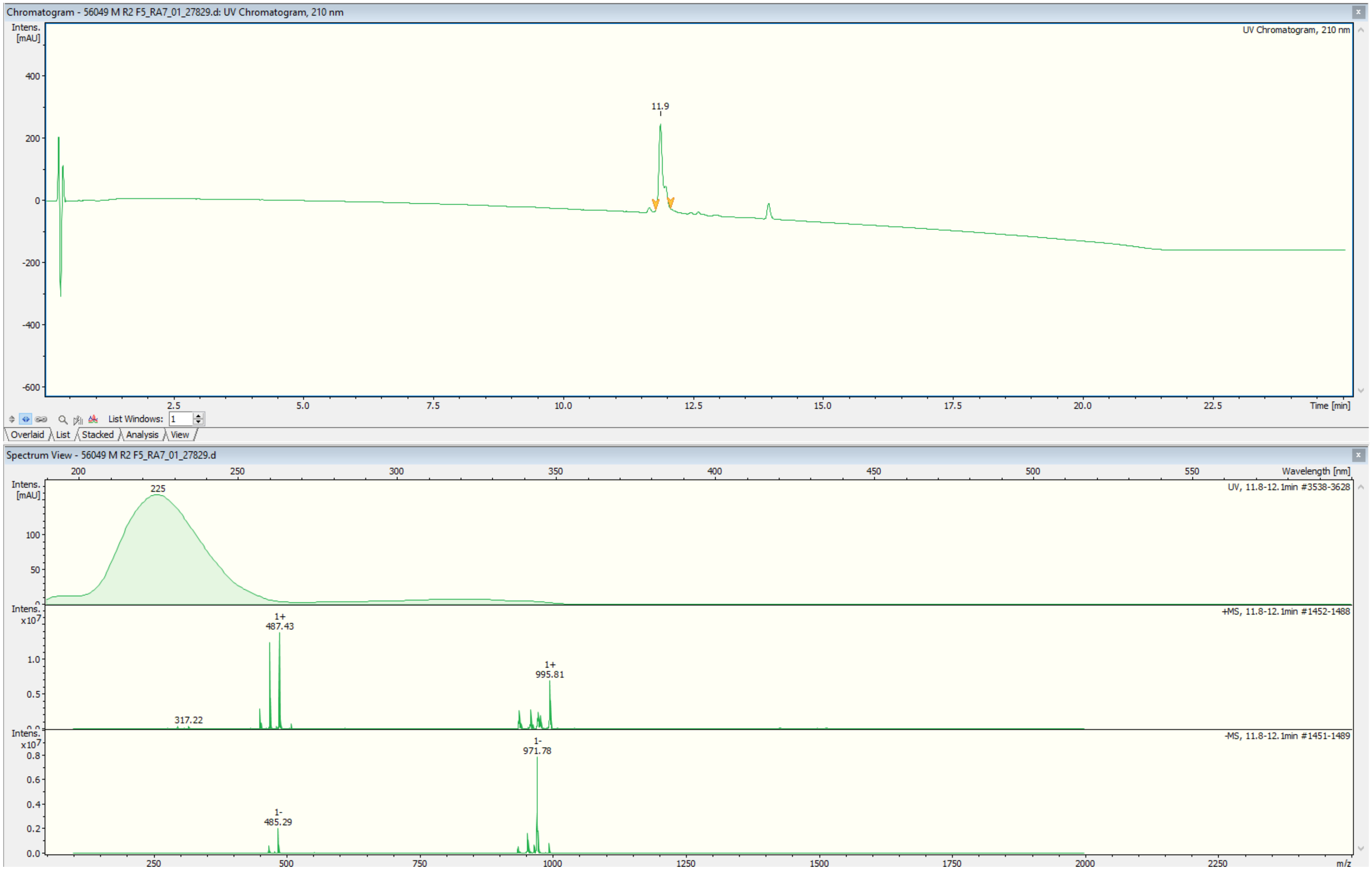


**Figure S1**: ESIMS data for tumulosic acid (**8**).

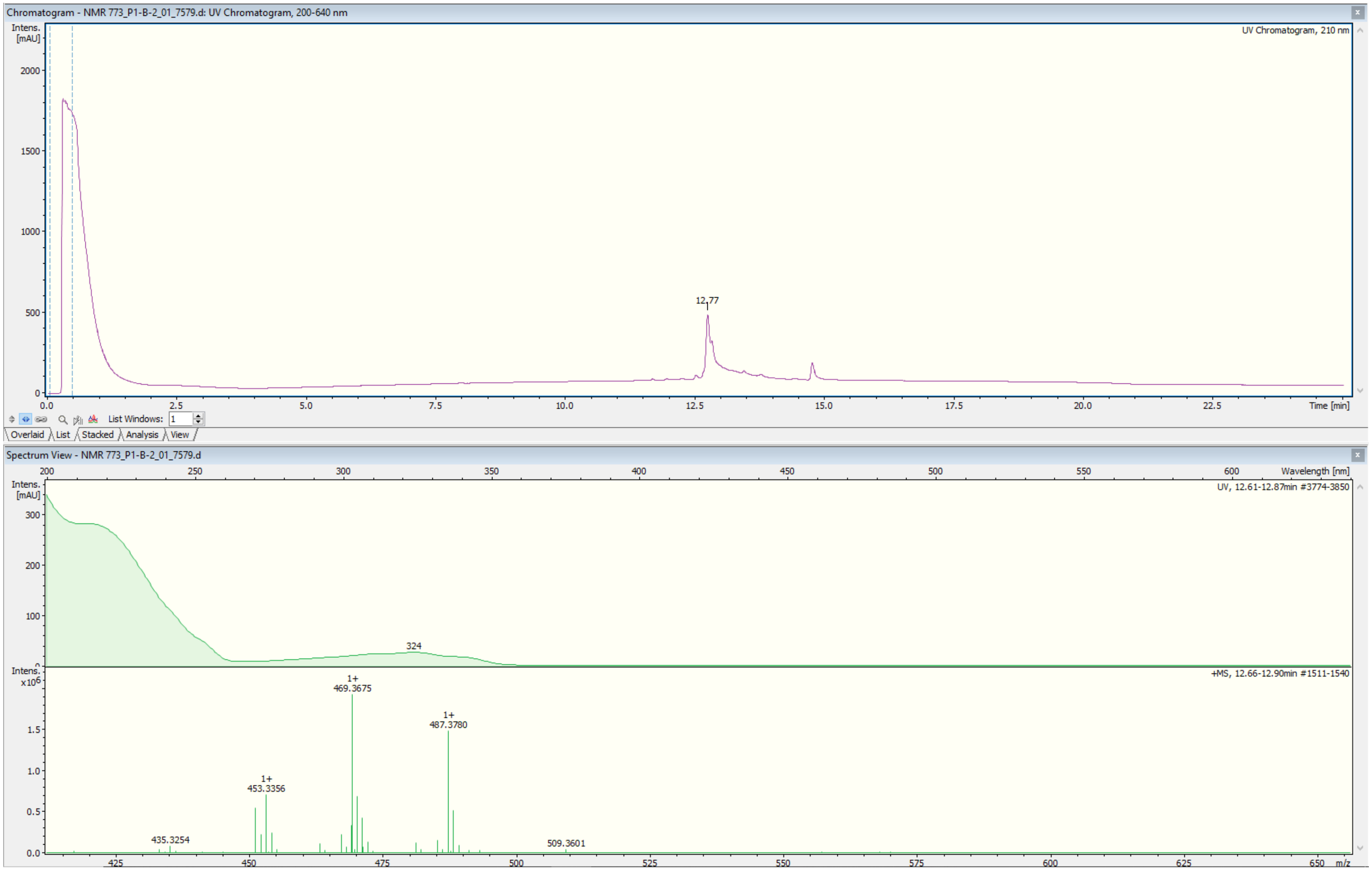


**Figure S2**: HR-ESIMS data for tumulosic acid (**8**).

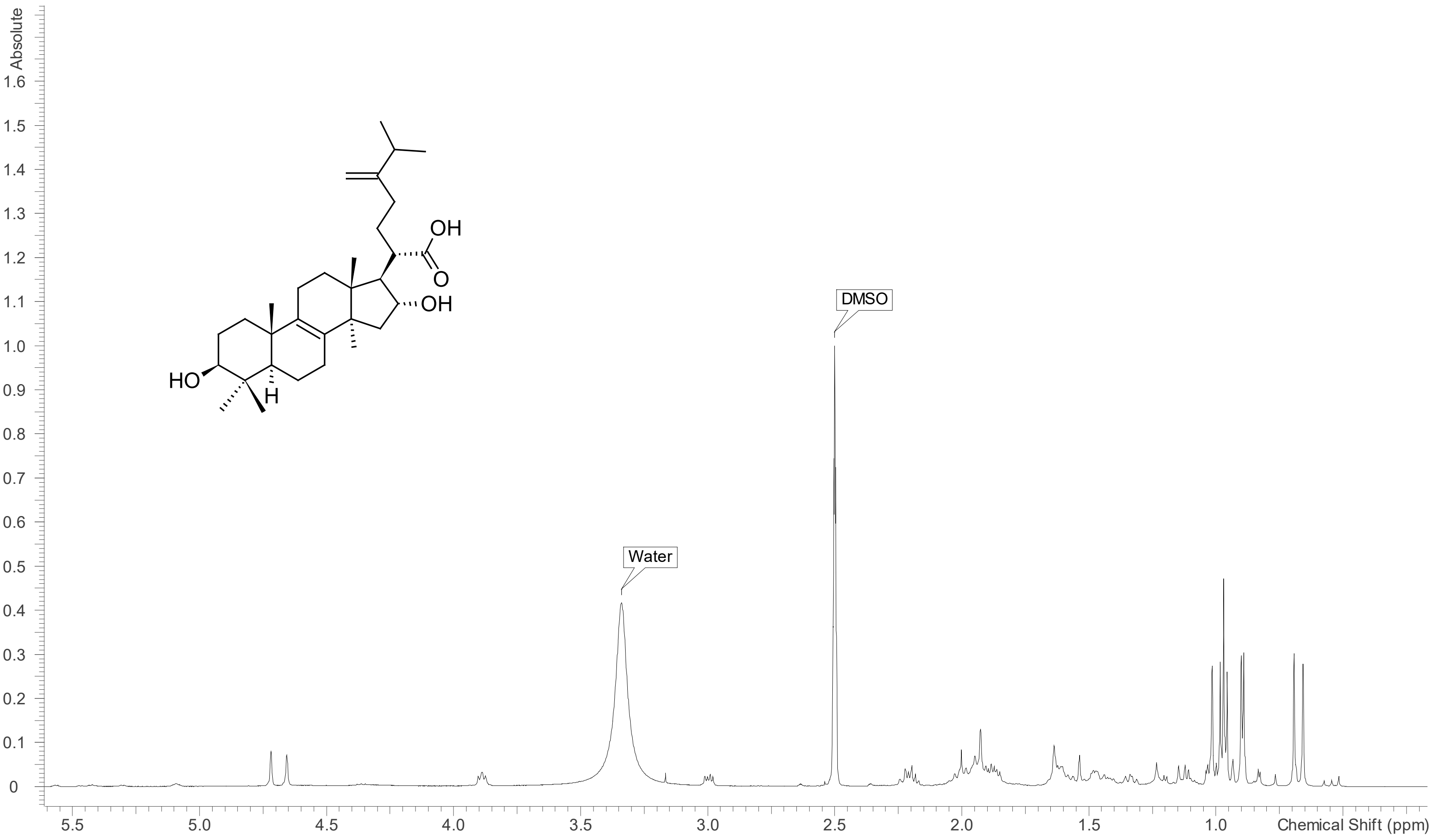


**Figure S3**: $^{1}$H NMR spectrum (DMSO-$d_6$, 700 MHz) of tumulosic acid (**8**).

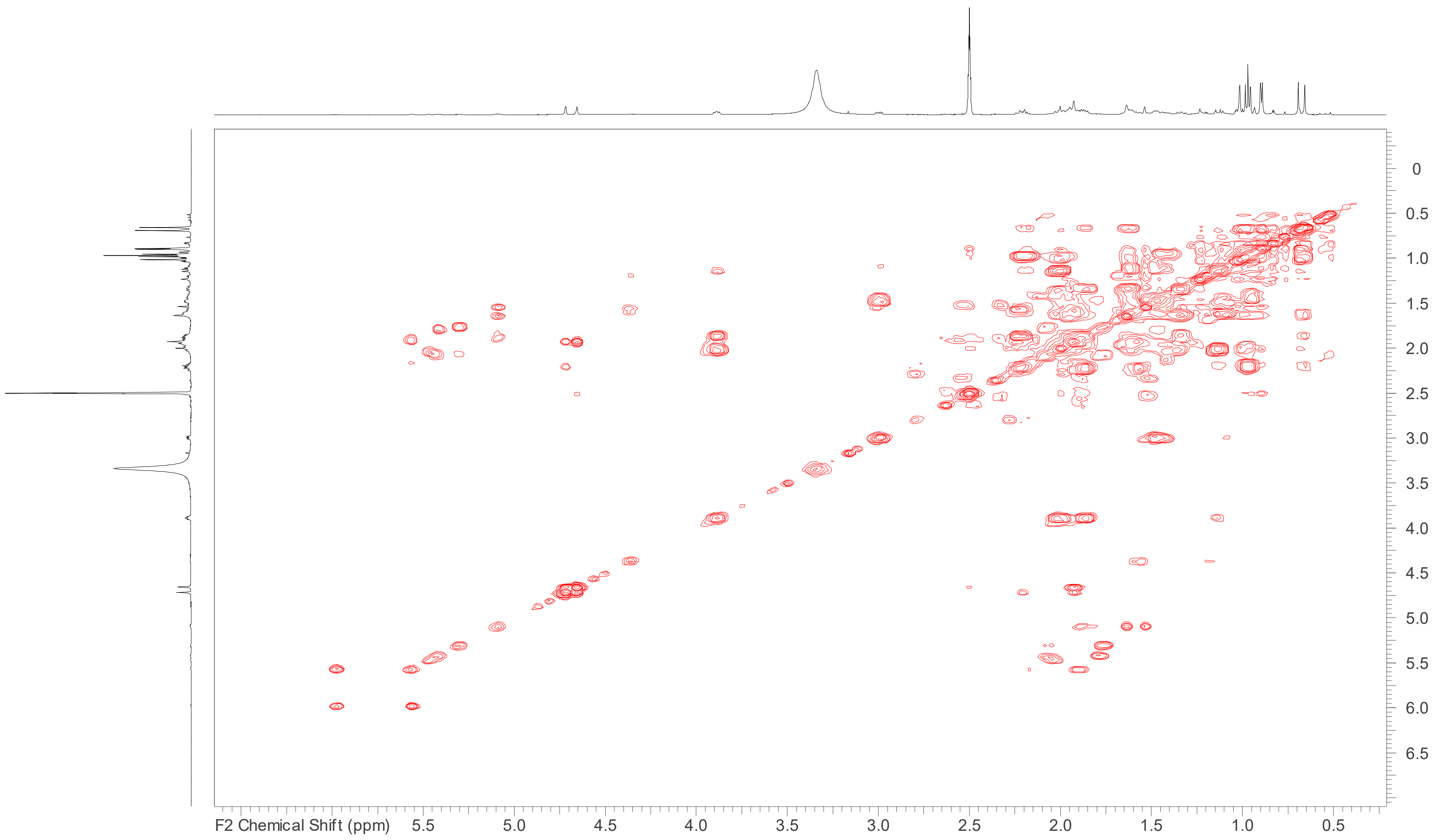


**Figure S4**: COSY spectrum (DMSO- $d_{6,}$ 700 MHz) of tumulosic acid (**8**).

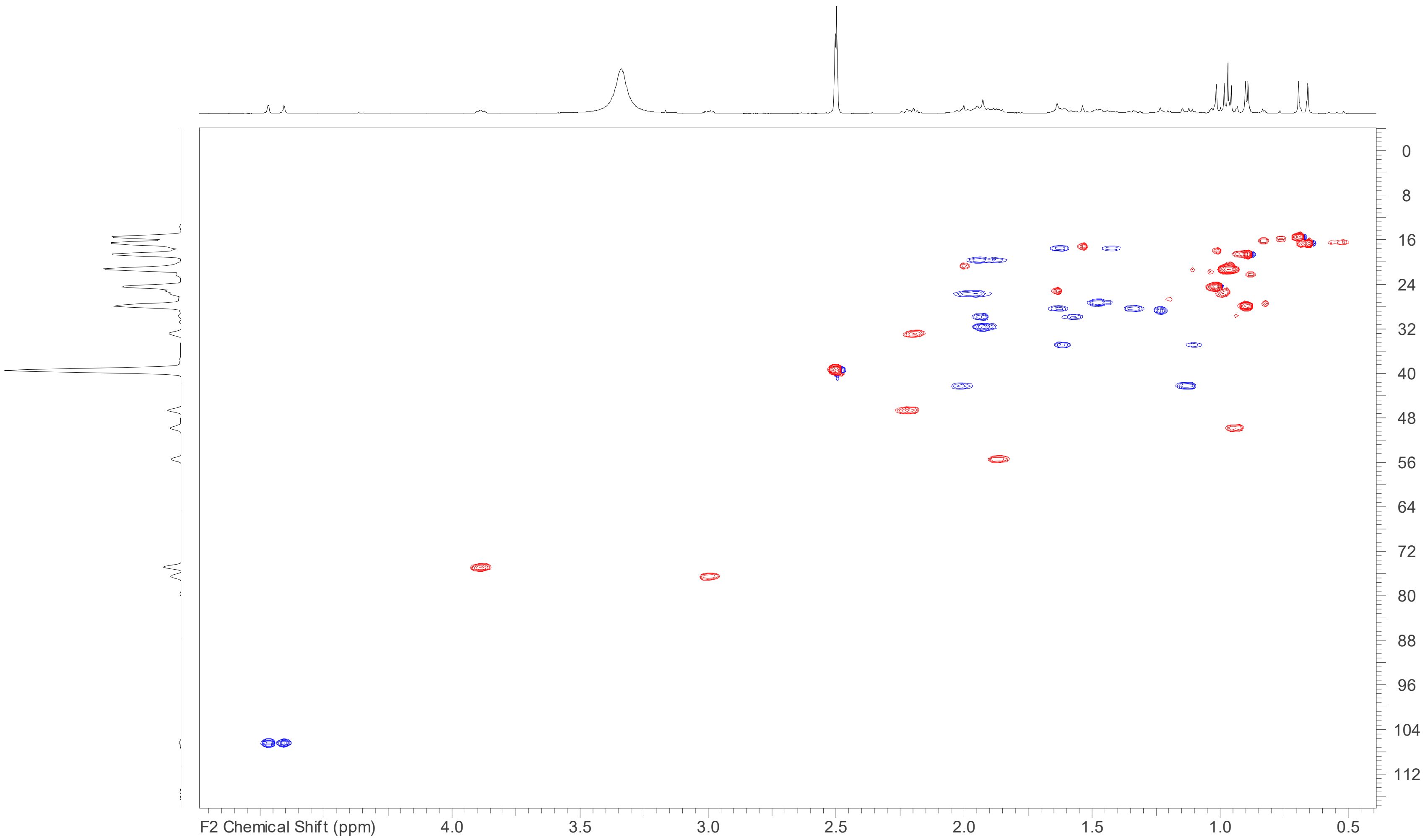


**Figure S5**: HSQC spectrum (DMSO- $d_6$, 700 MHz) of tumulosic acid (**8**).

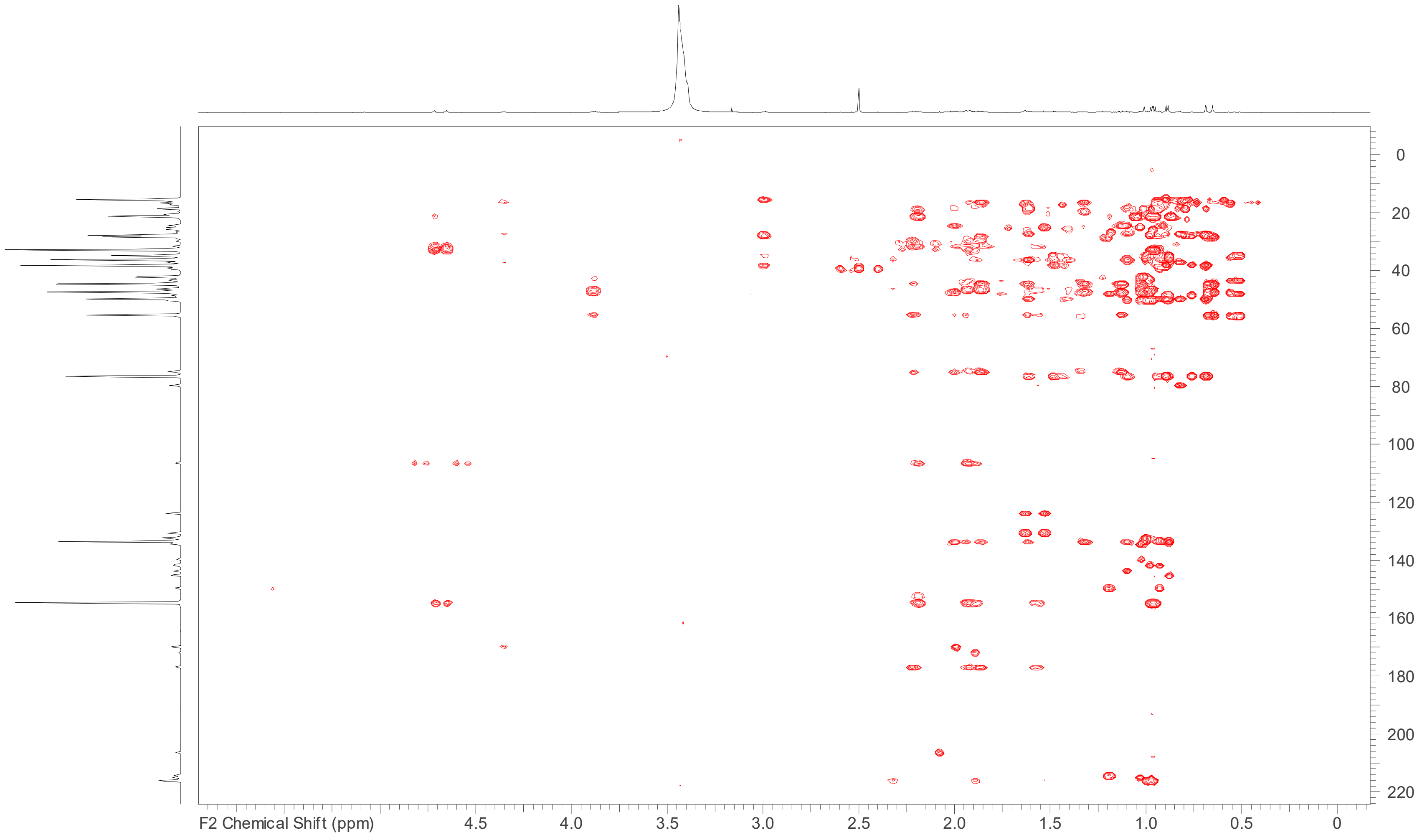


**Figure S6**: HMBC spectrum (DMSO-$d_6$, 700 MHz) of tumulosic acid (**8**).

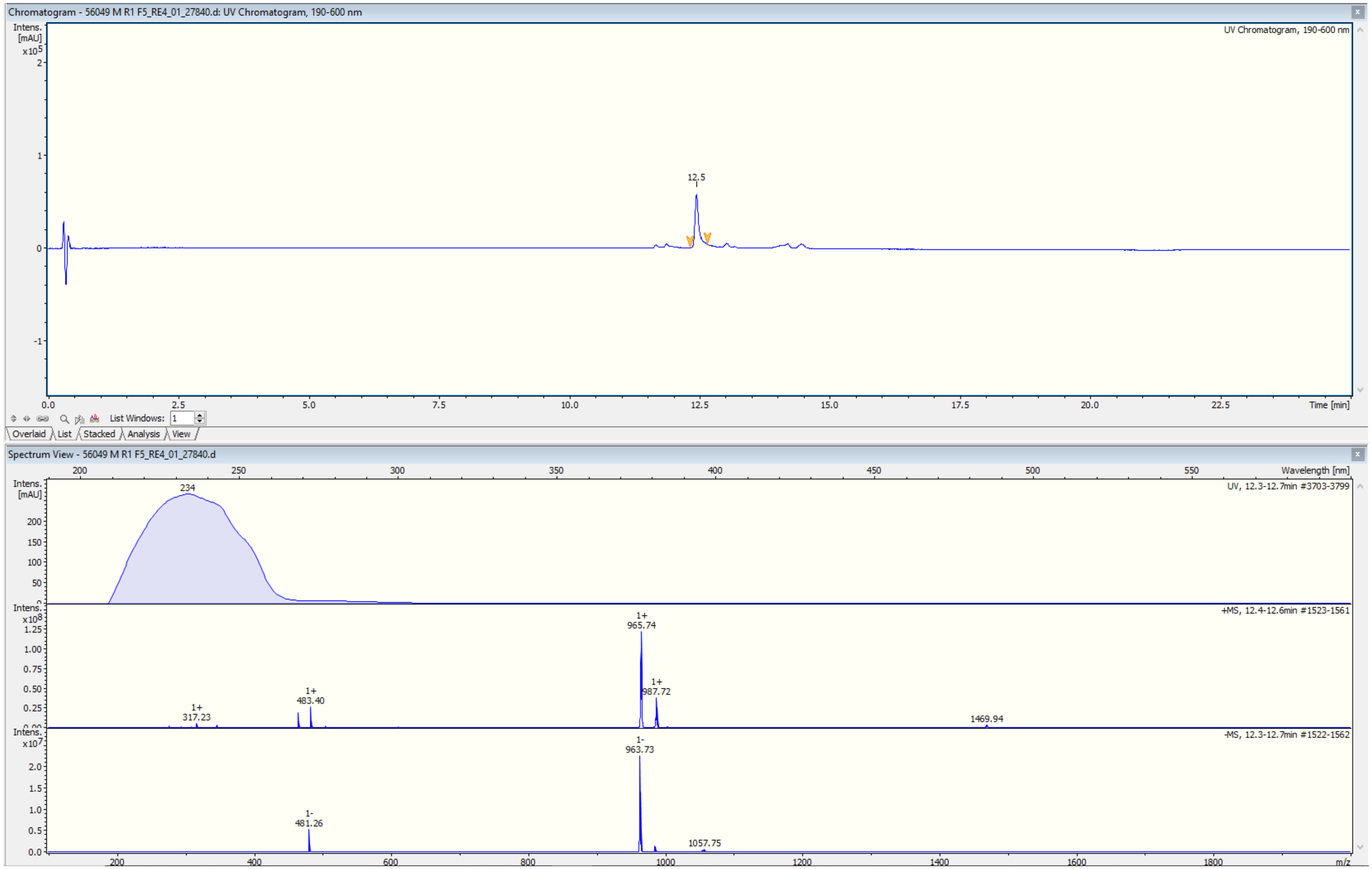


**Figure S7**: ESIMS data for polyporenic acid C (**9**).

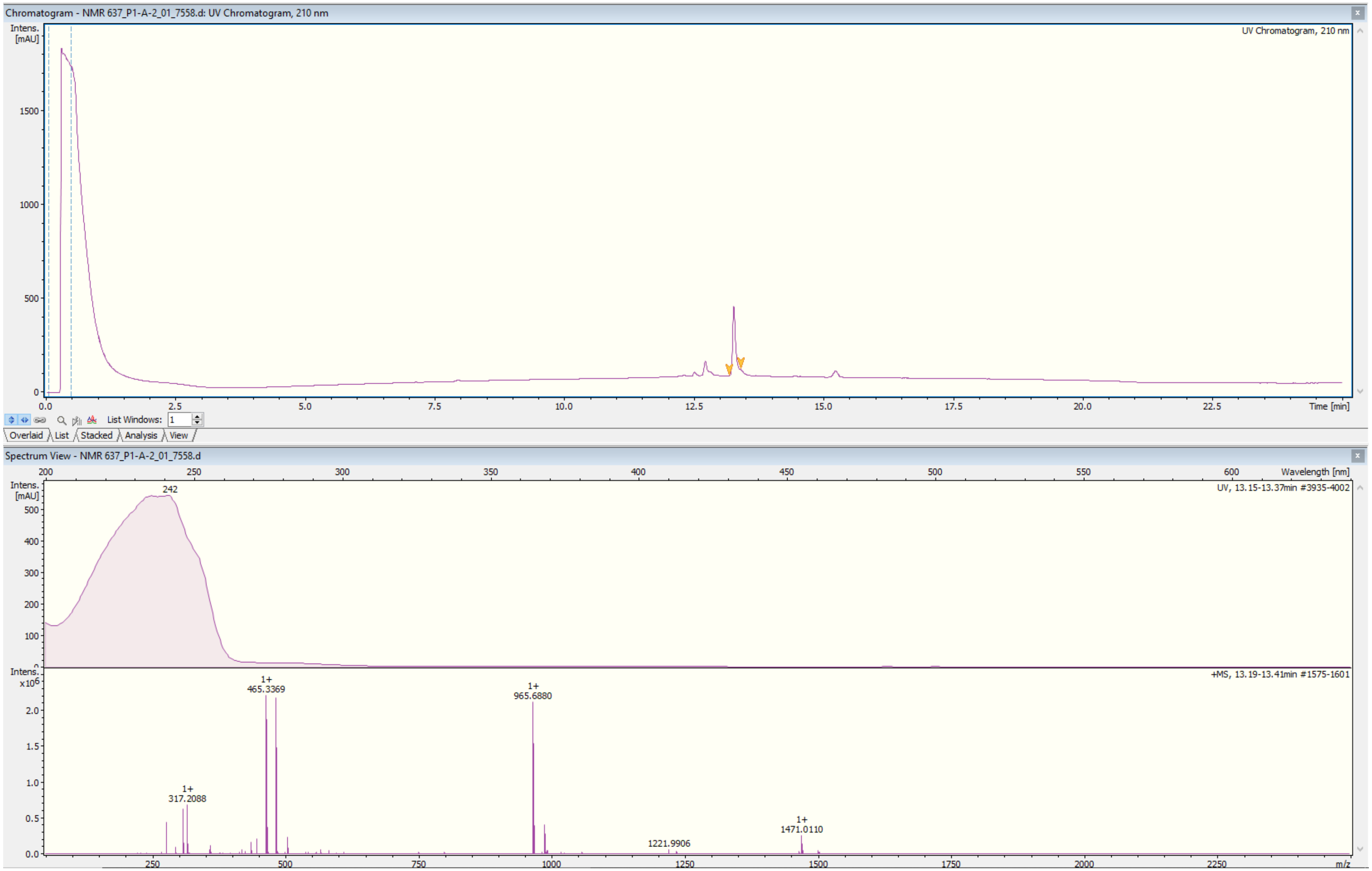


**Figure S8**: HR-ESIMS data for polyporenic acid C (**9**).

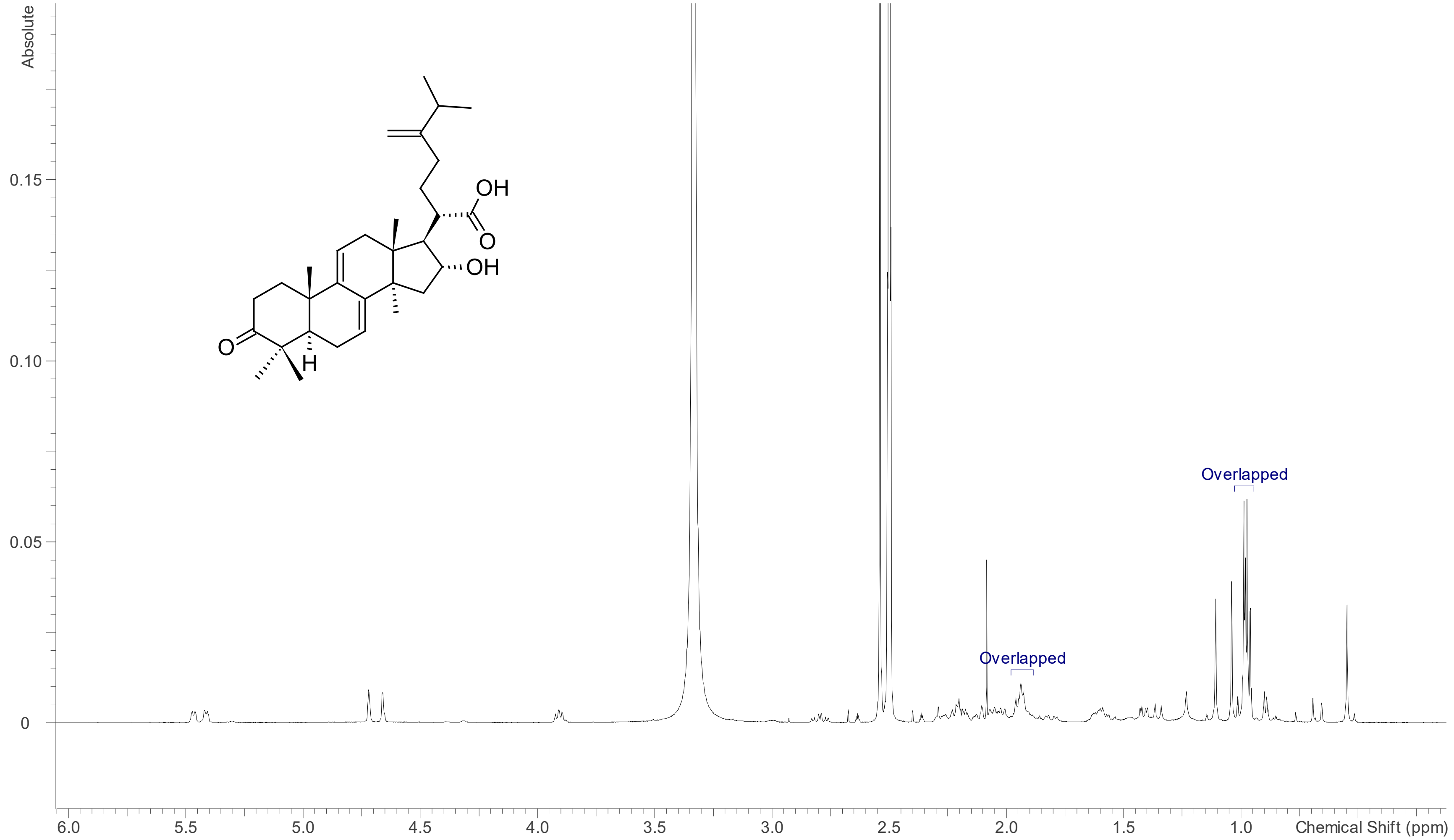


**Figure S9**: [1]H NMR spectrum (DMSO-$d_6$, 700 MHz) of polyporenic acid C (**9**).

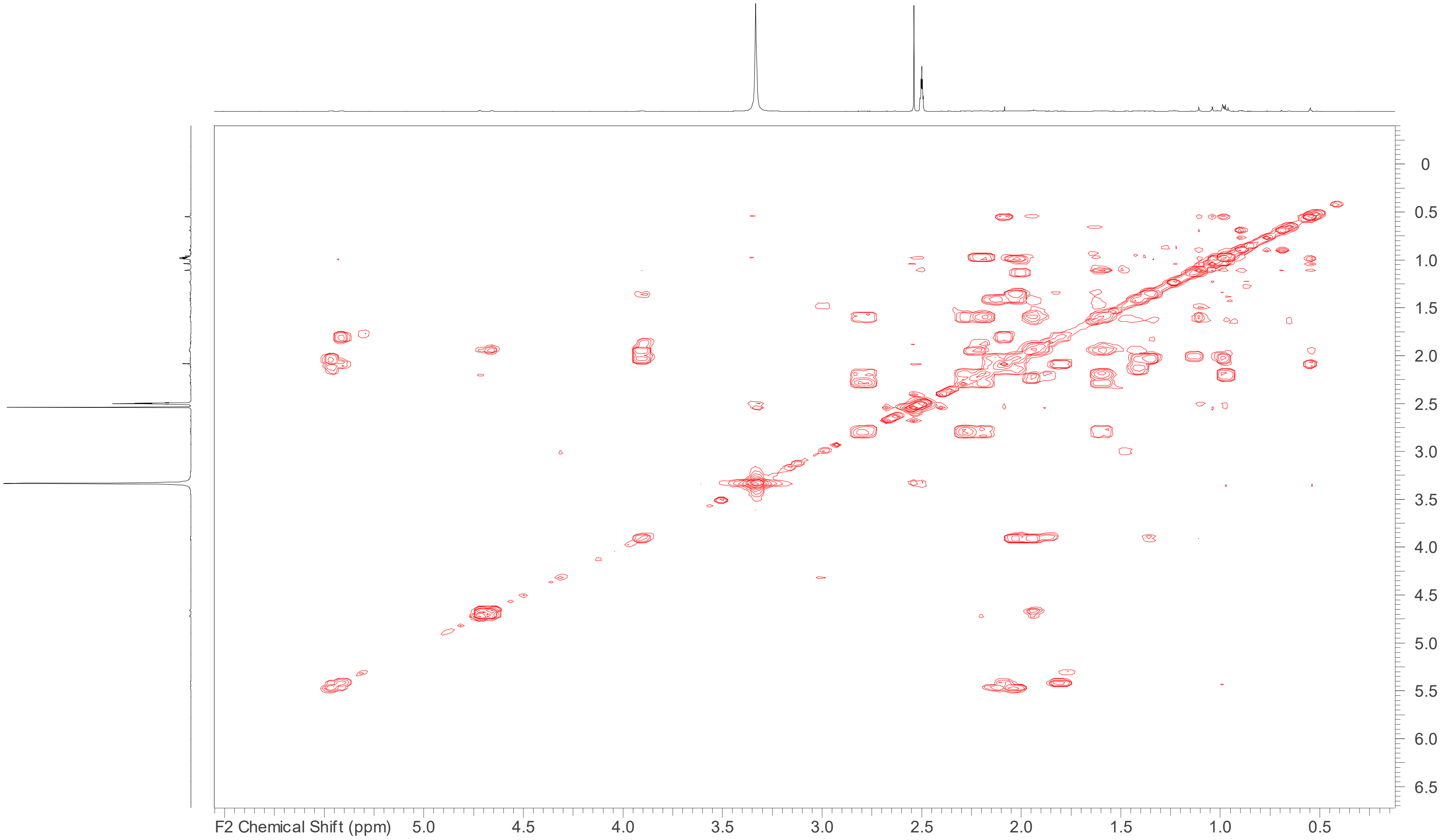


**Figure S10**: COSY spectrum (DMSO-$d_6$, 700 MHz) of polyporenic acid C (**9**).

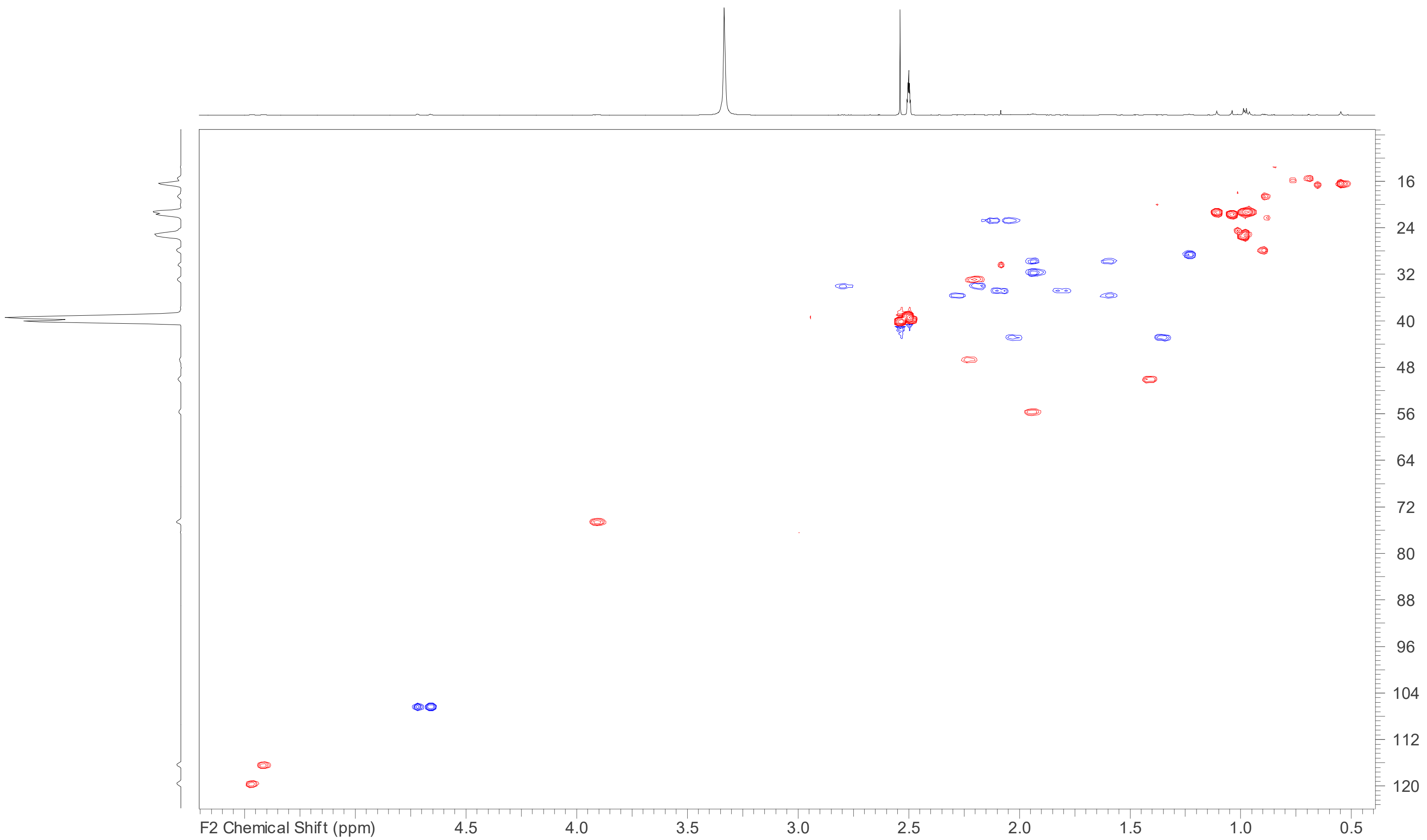


**Figure S11**: HSQC spectrum (DMSO-$d_6$, 700 MHz) of polyporenic acid C (**9**).

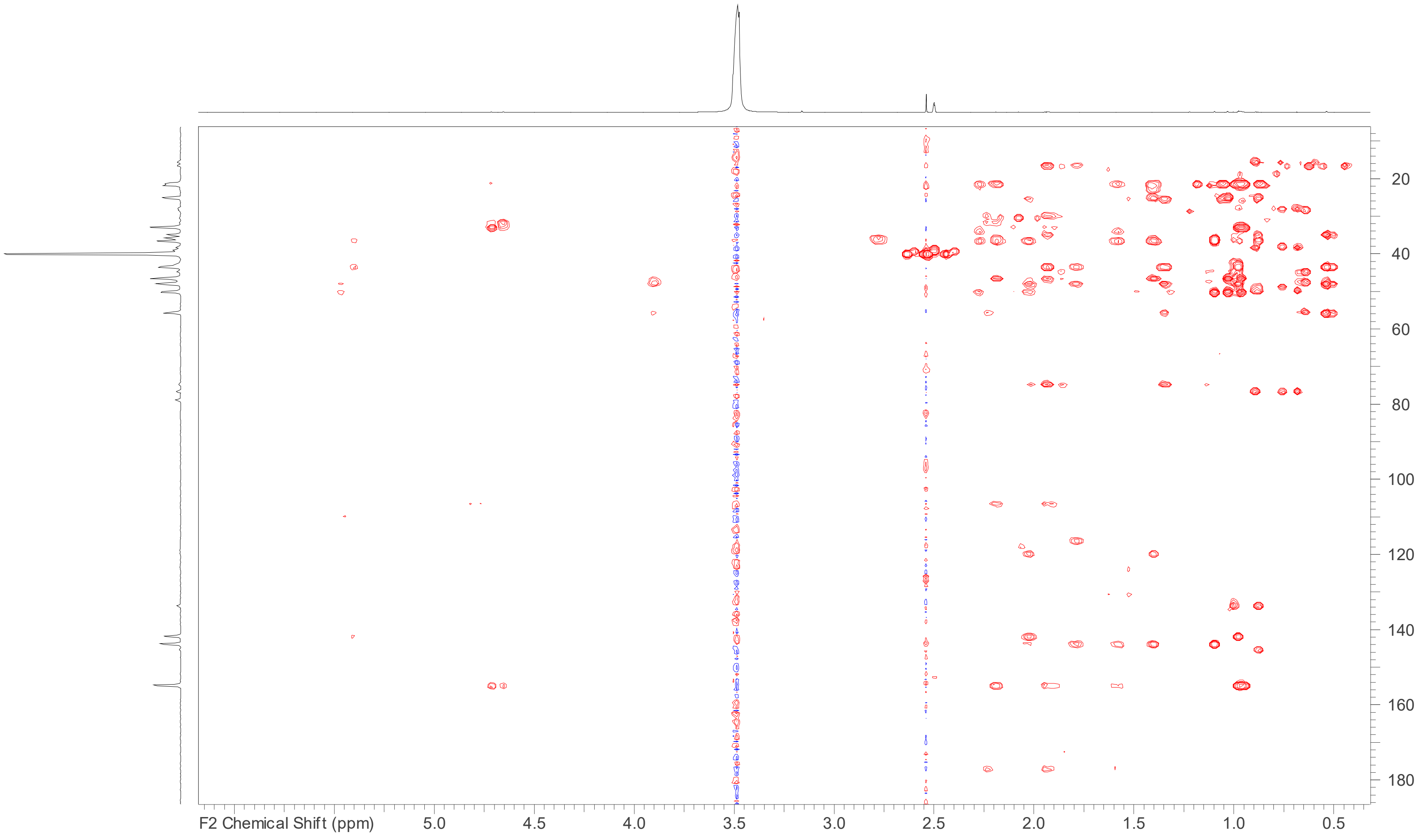


**Figure S12**: HMBC spectrum (DMSO-$d_6$, 700 MHz) of polyporenic acid C (**9**).

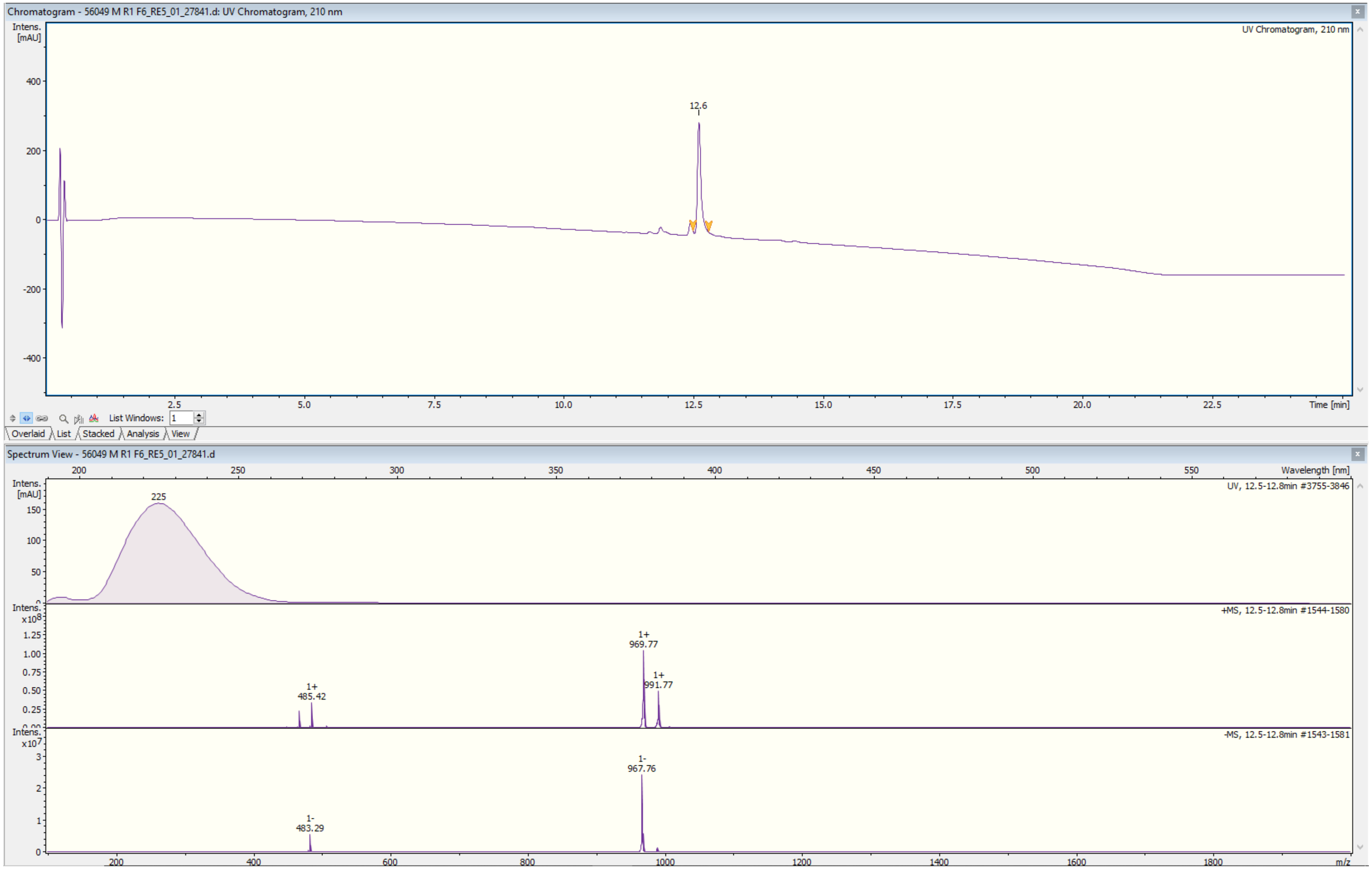


**Figure S13**: ESIMS data for 16α-hydroxyeburiconic acid (**10**).

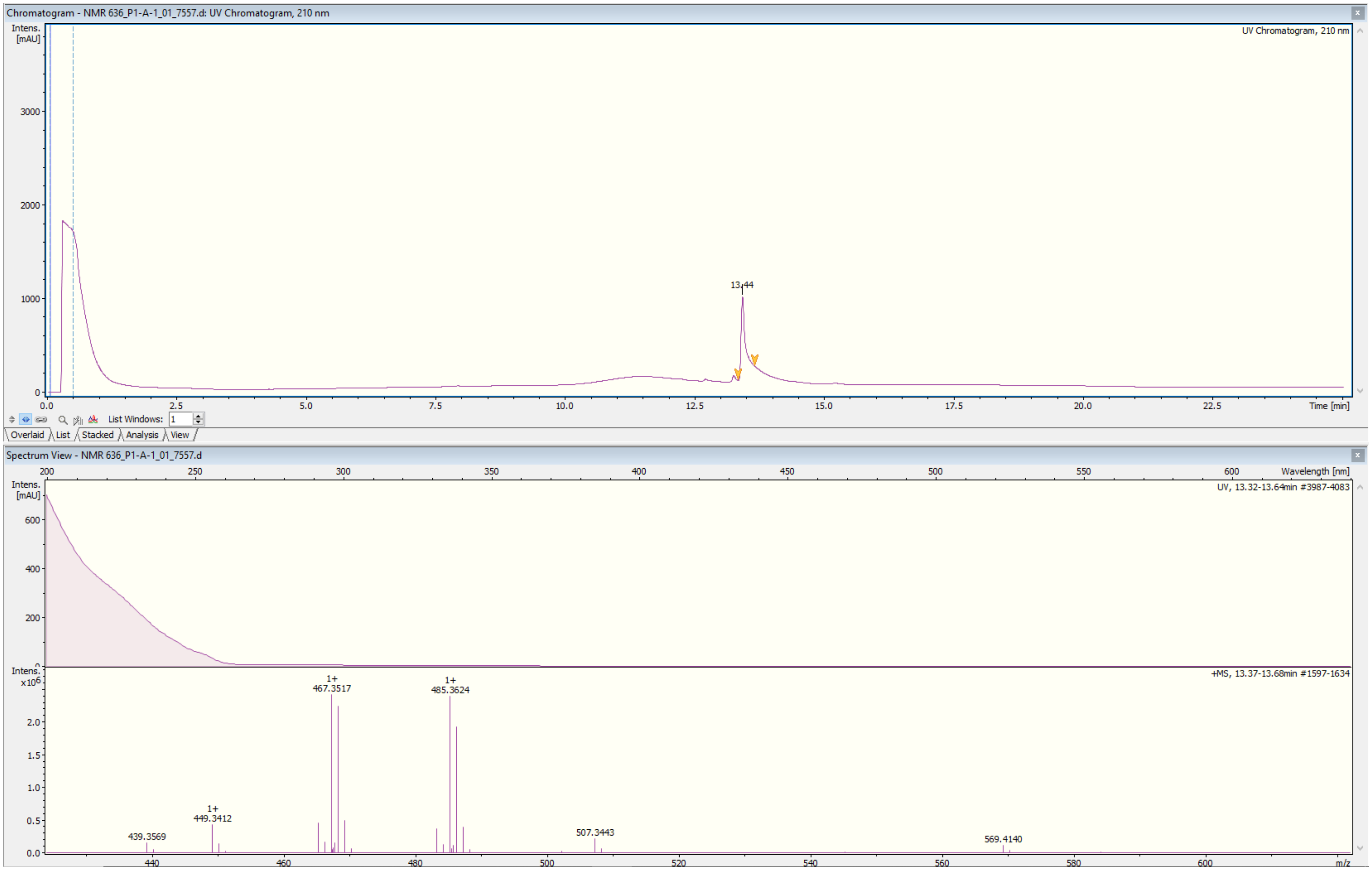


**Figure S14**: HR-ESIMS data for 16α-hydroxyeburiconic acid (**10**).

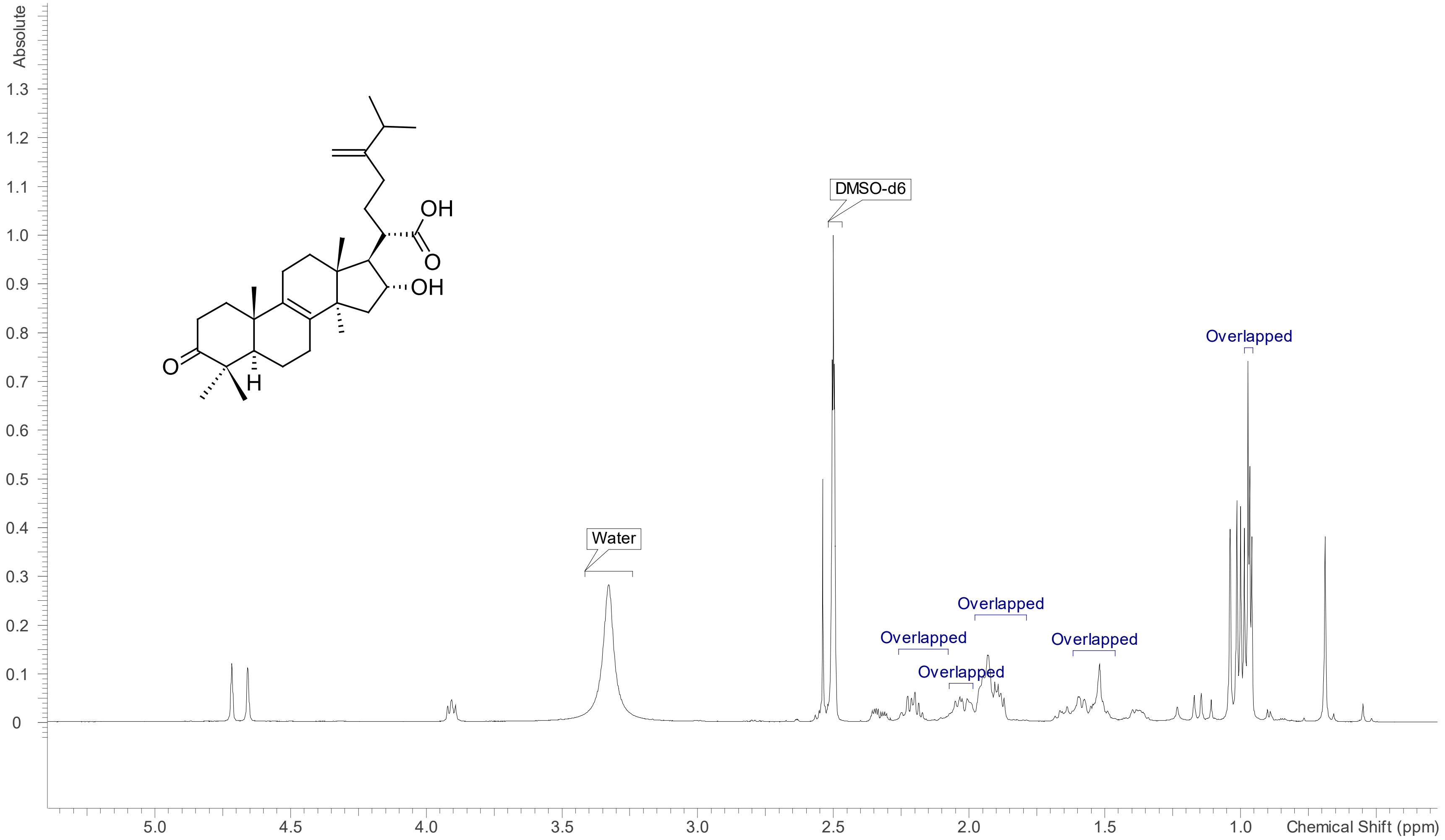


**Figure S15**: [1]H NMR spectrum (DMSO-$d_6$, 700 MHz) of 16α-hydroxyeburiconic acid (**10**).

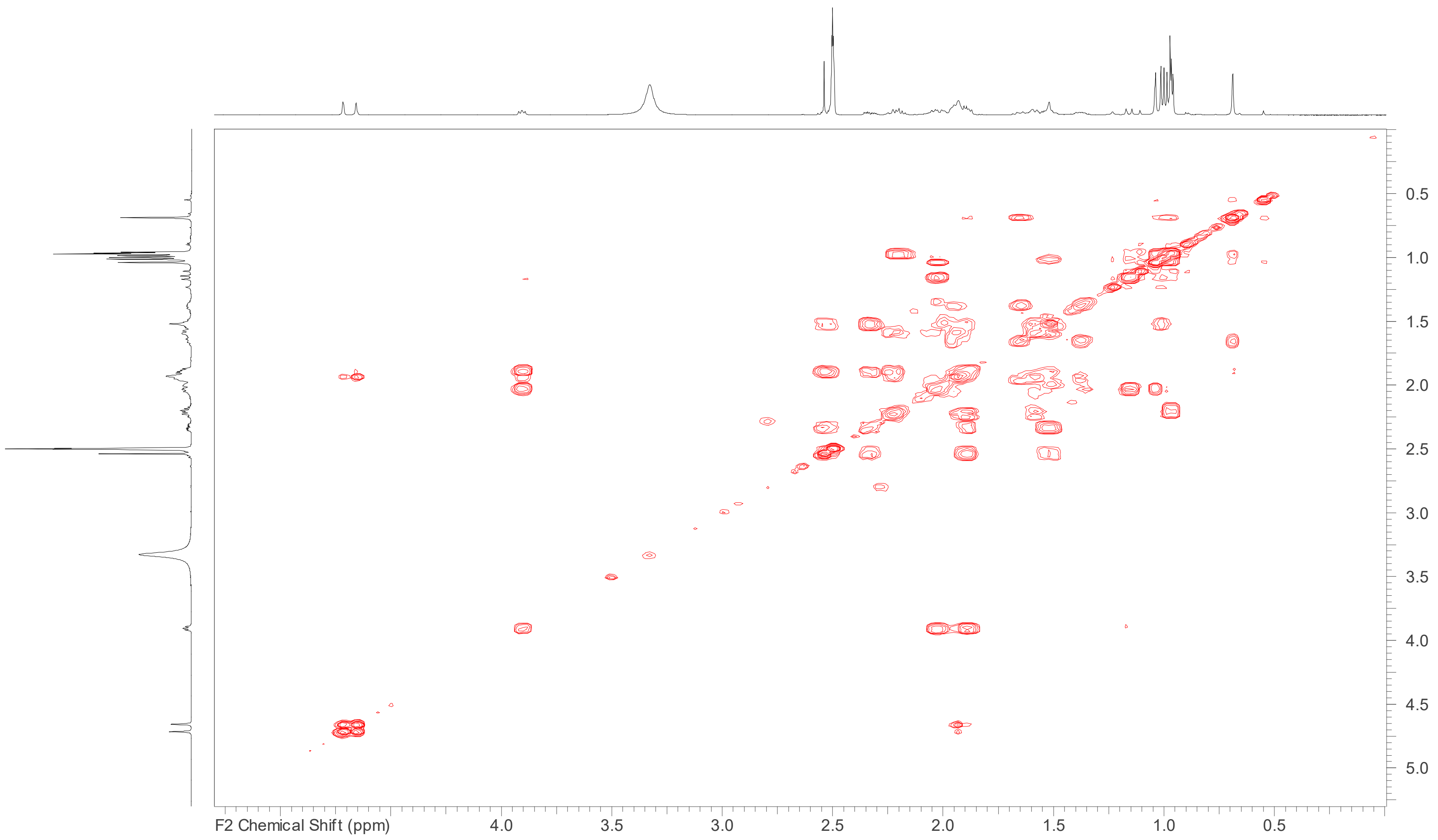


**Figure S16**: COSY spectrum (DMSO- $d_6$, 700 MHz) of 16α-hydroxyeburiconic acid (**10**).

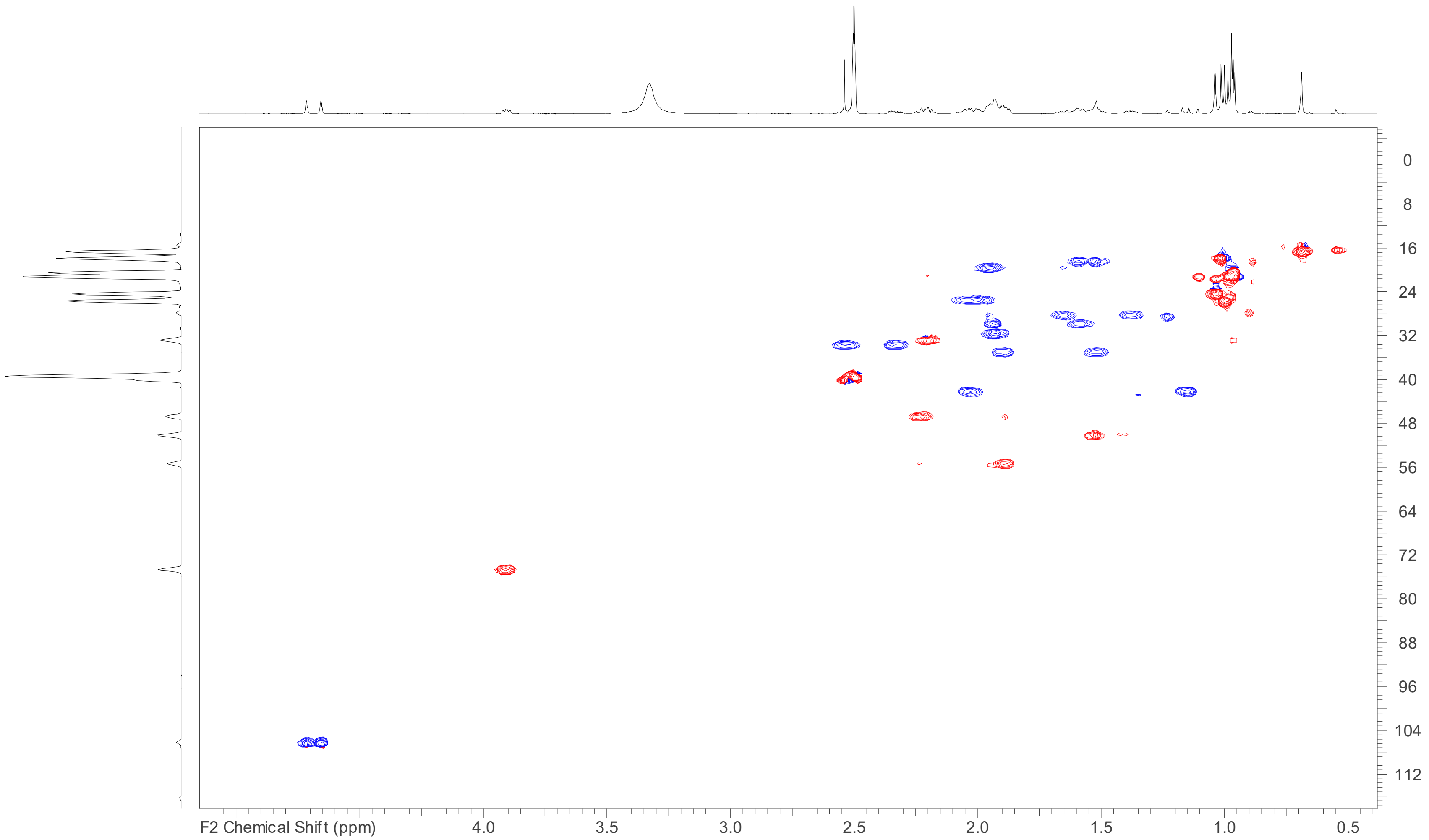


**Figure S17**: HSQC spectrum (DMSO-$d_6$, 700 MHz) of 16α-hydroxyeburiconic acid (**10**).

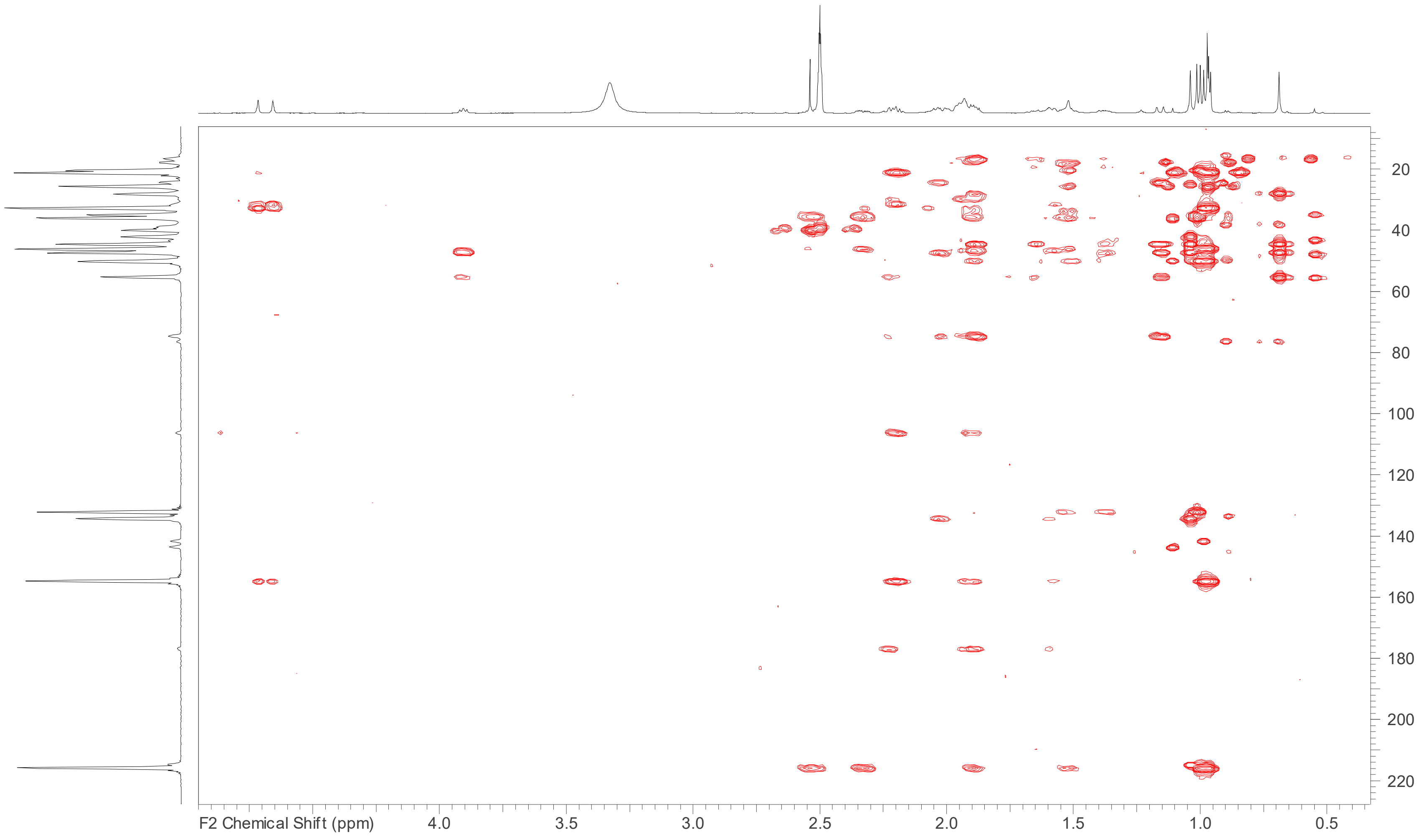


**Figure S18**: HMBC spectrum (DMSO-$d_6$, 700 MHz) of 16α-hydroxyeburiconic acid (**10**).

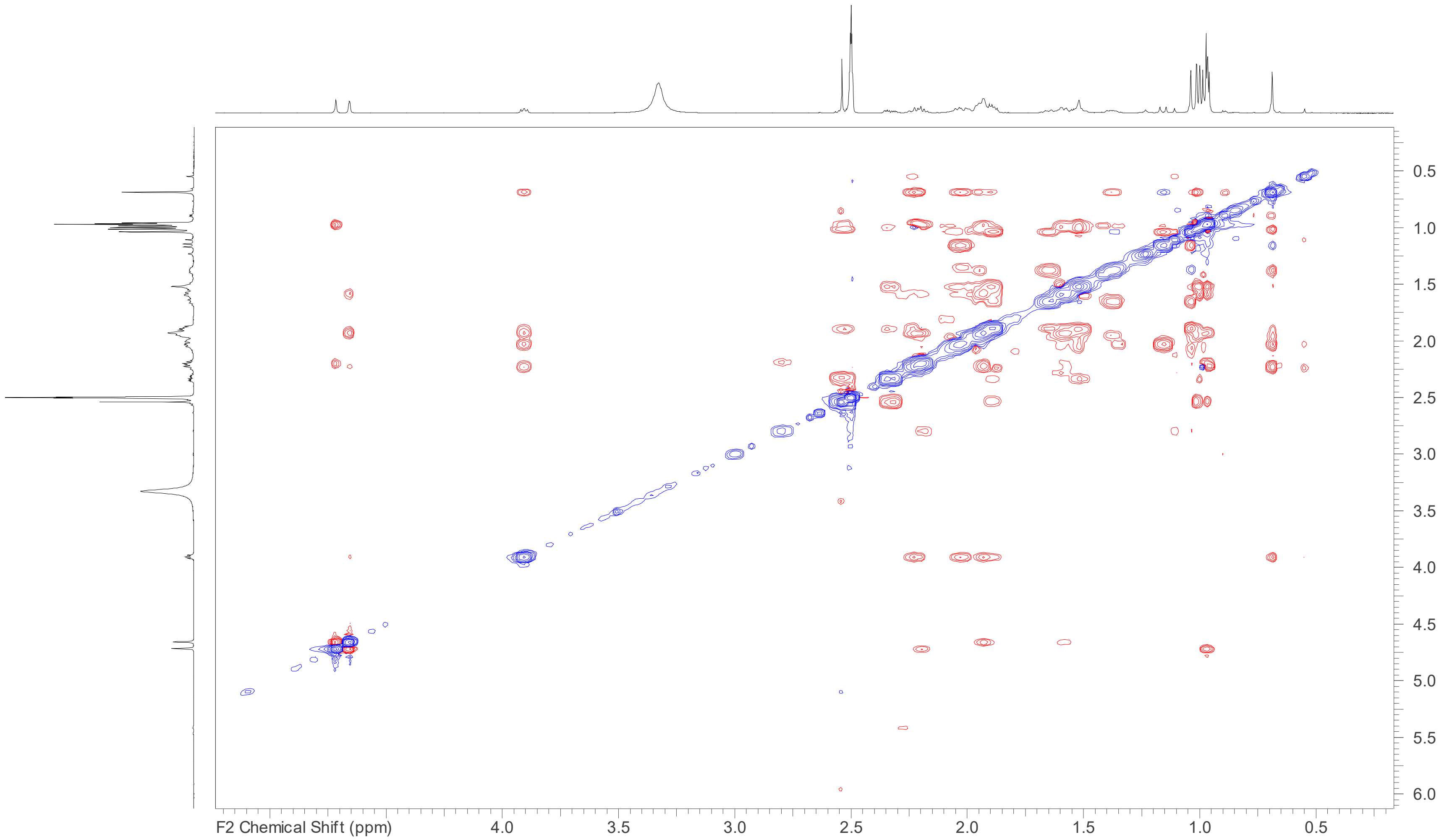


**Figure S19**: ROESY spectrum (DMSO-$d_6$, 700 MHz) of 16α-hydroxyeburiconic acid (**10**).

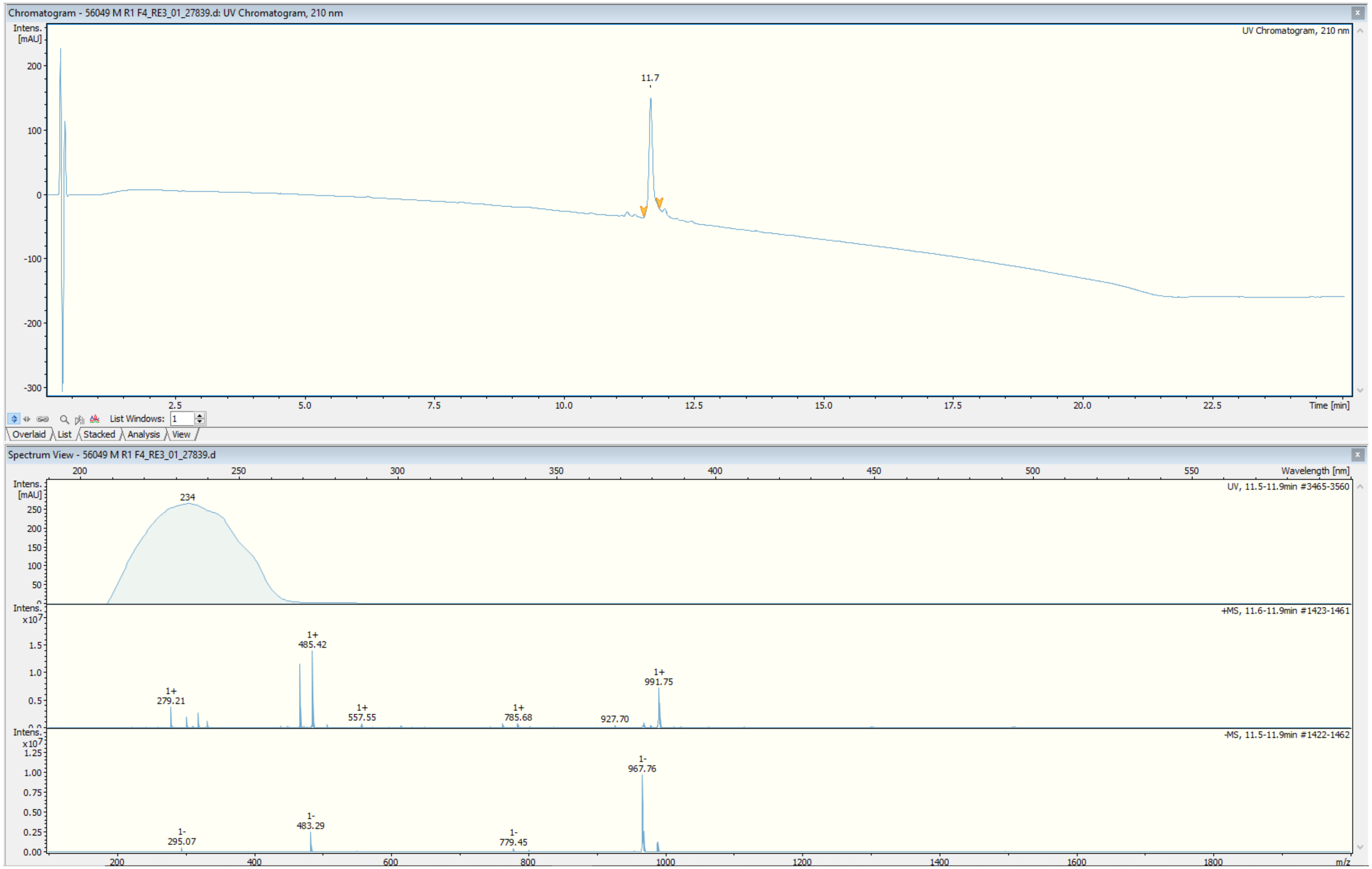


**Figure S20**: ESIMS data for dehydrotumulosic acid (**11**)

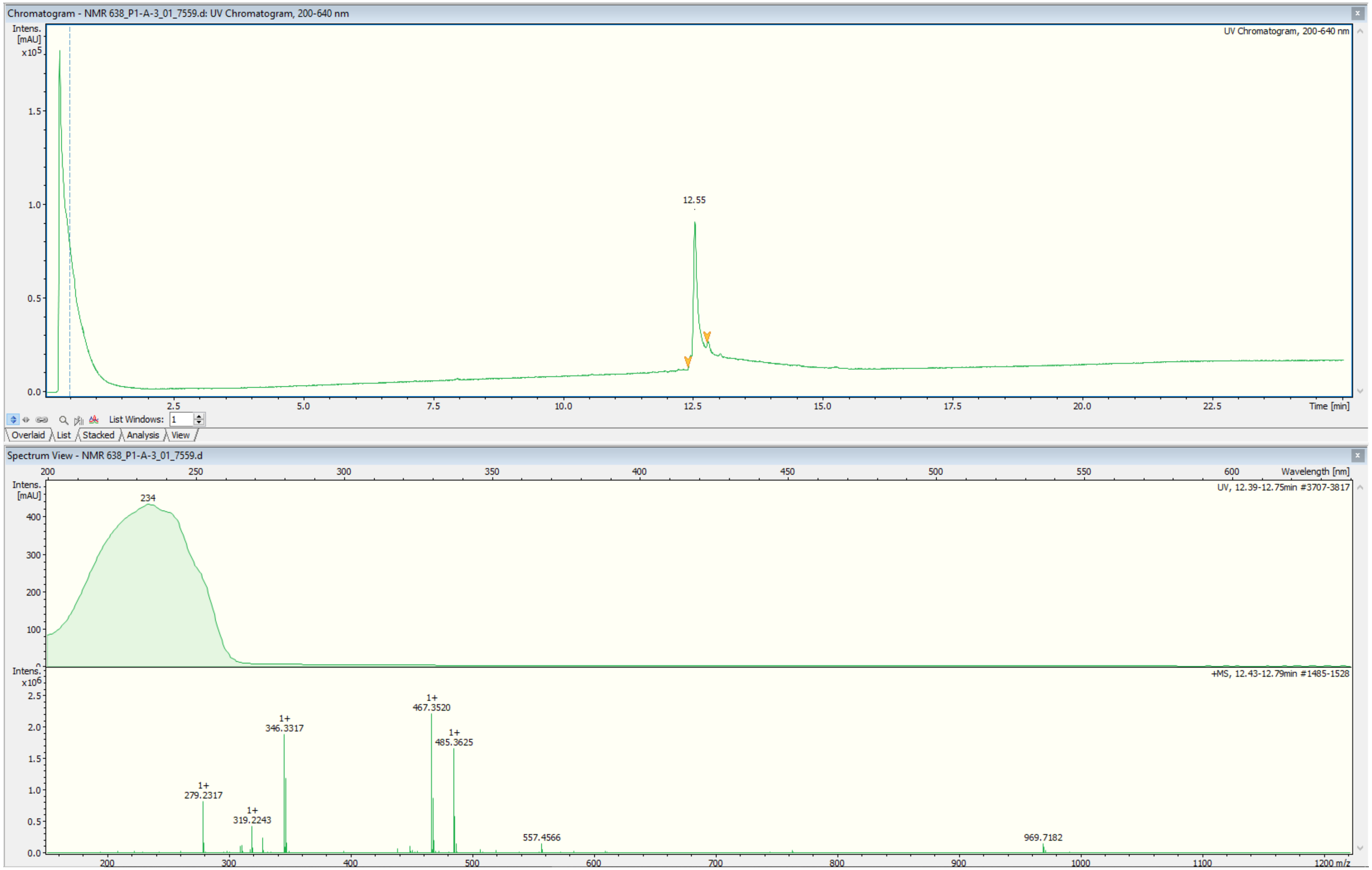


**Figure S21**: HR-ESIMS data for dehydrotumulosic acid (**11**).

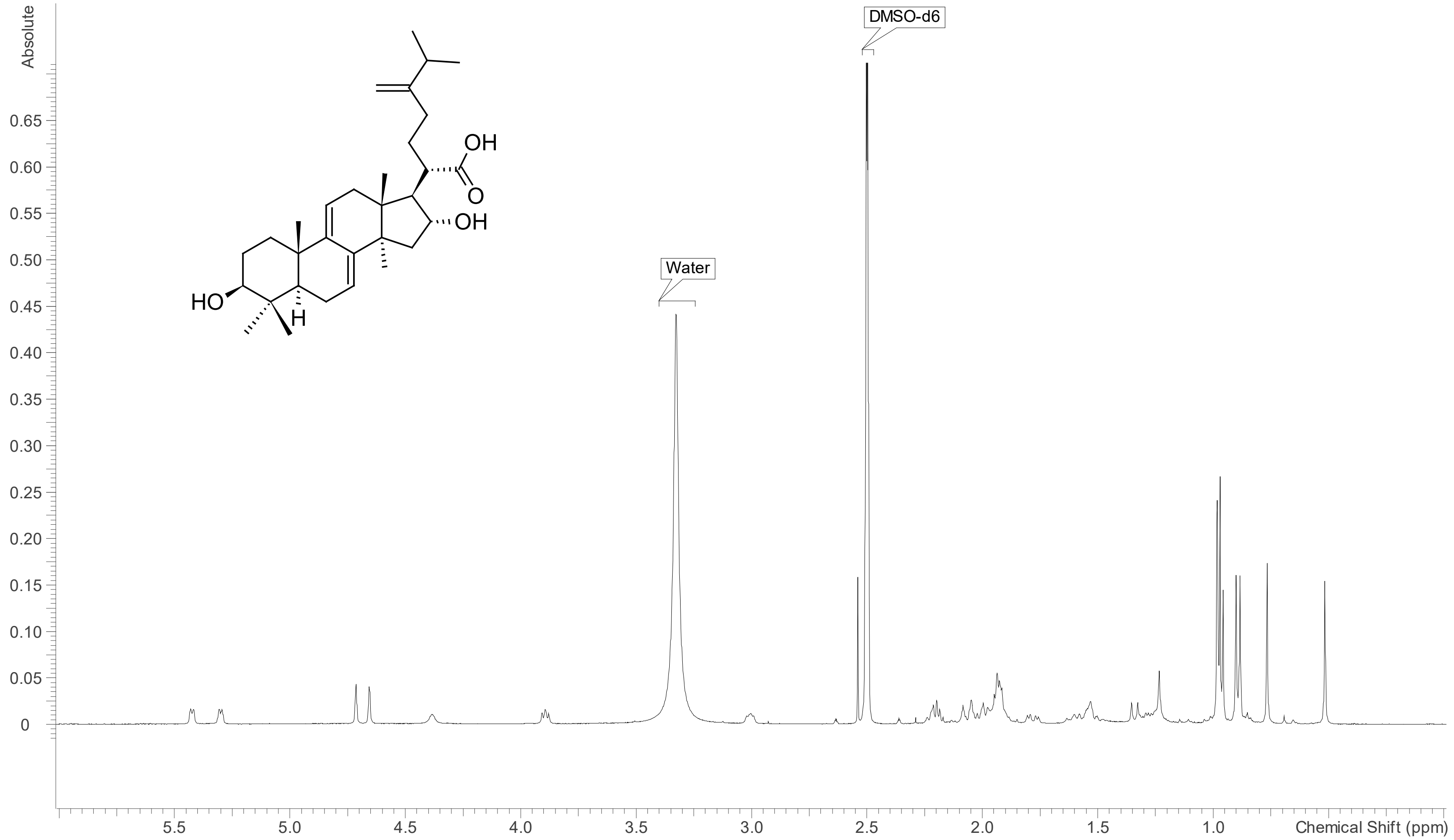


**Figure S22**: $^{1}$H NMR spectrum (DMSO-$d_6$, 700 MHz) of dehydrotumulosic acid (**11**).

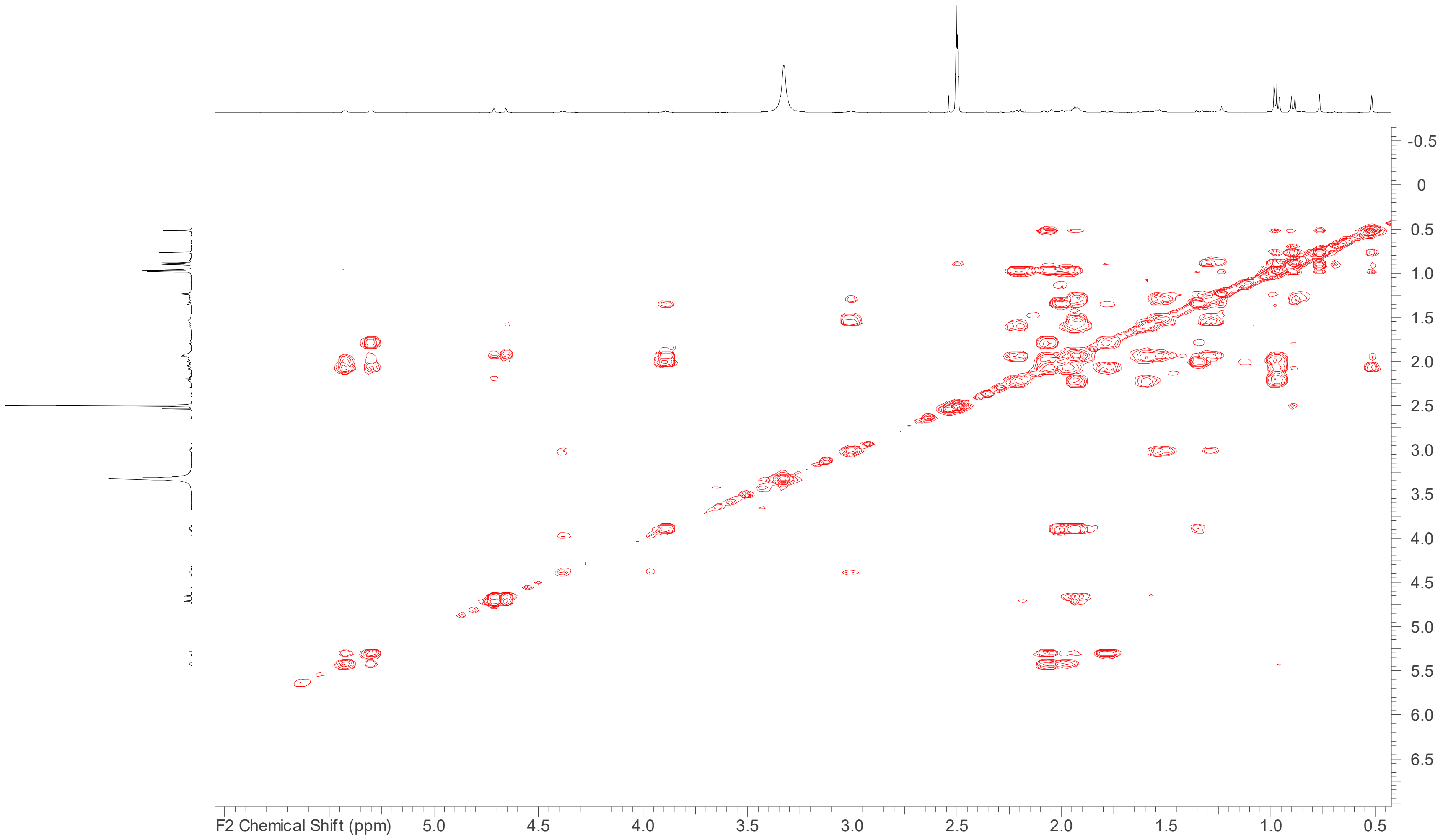


**Figure S23**: COSY spectrum (DMSO-$d_6$, 700 MHz) of dehydrotumulosic acid (**11**).

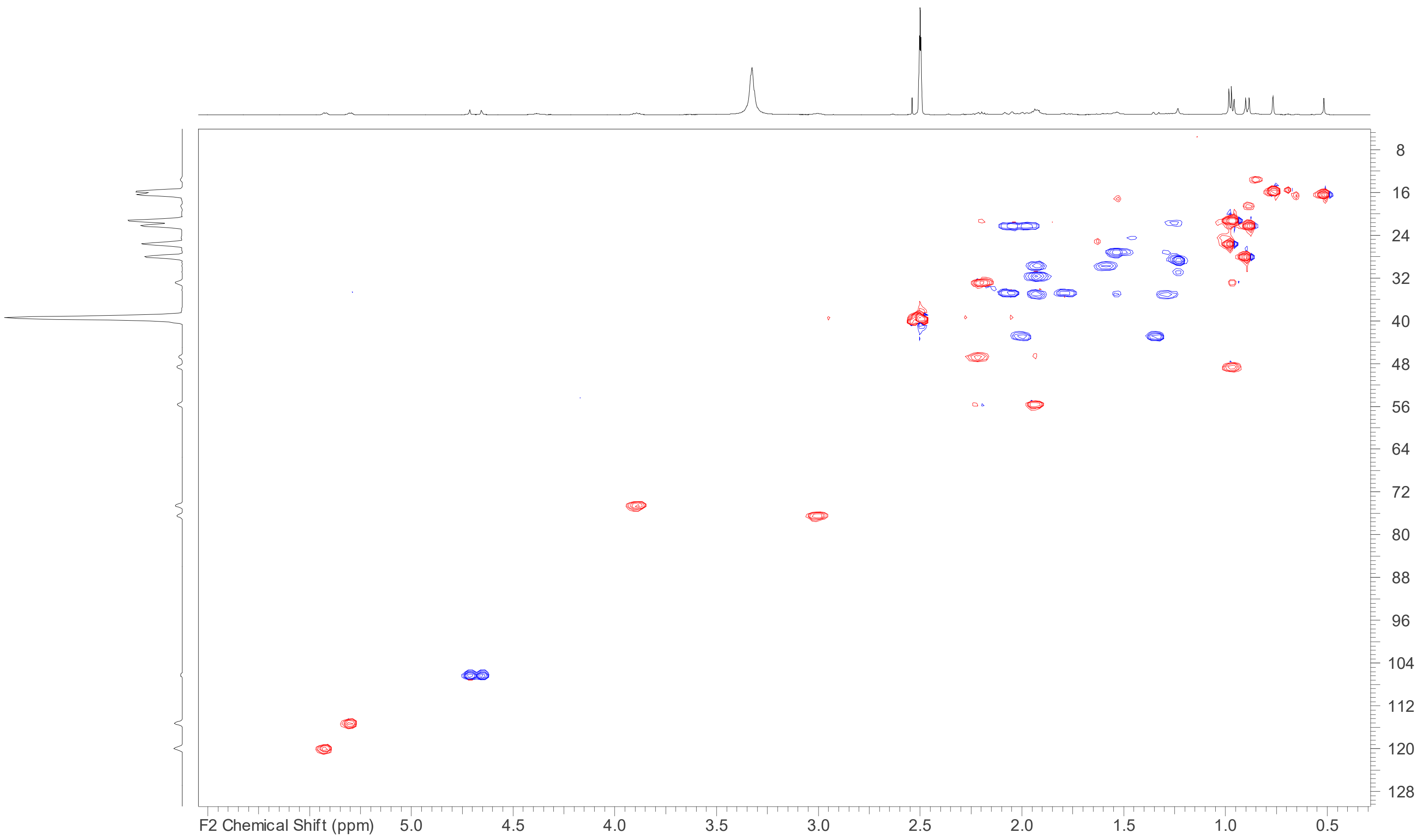


**Figure S24**: HSQC spectrum (DMSO-$d_6$, 700 MHz) of dehydrotumulosic acid (**11**).

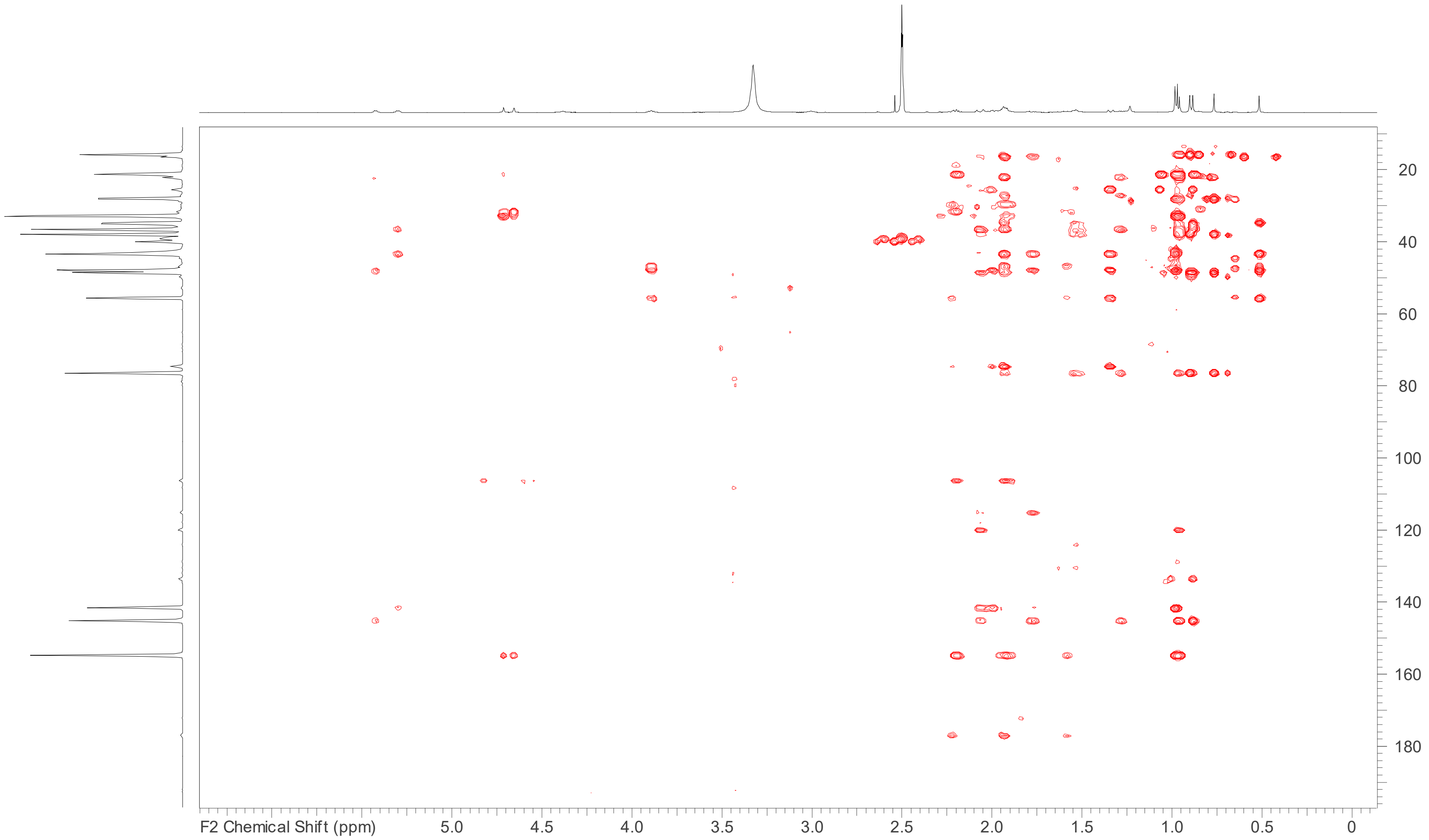


**Figure S25**: HMBC spectrum (DMSO-$d_6$, 700 MHz) of dehydrotumulosic acid (**11**).

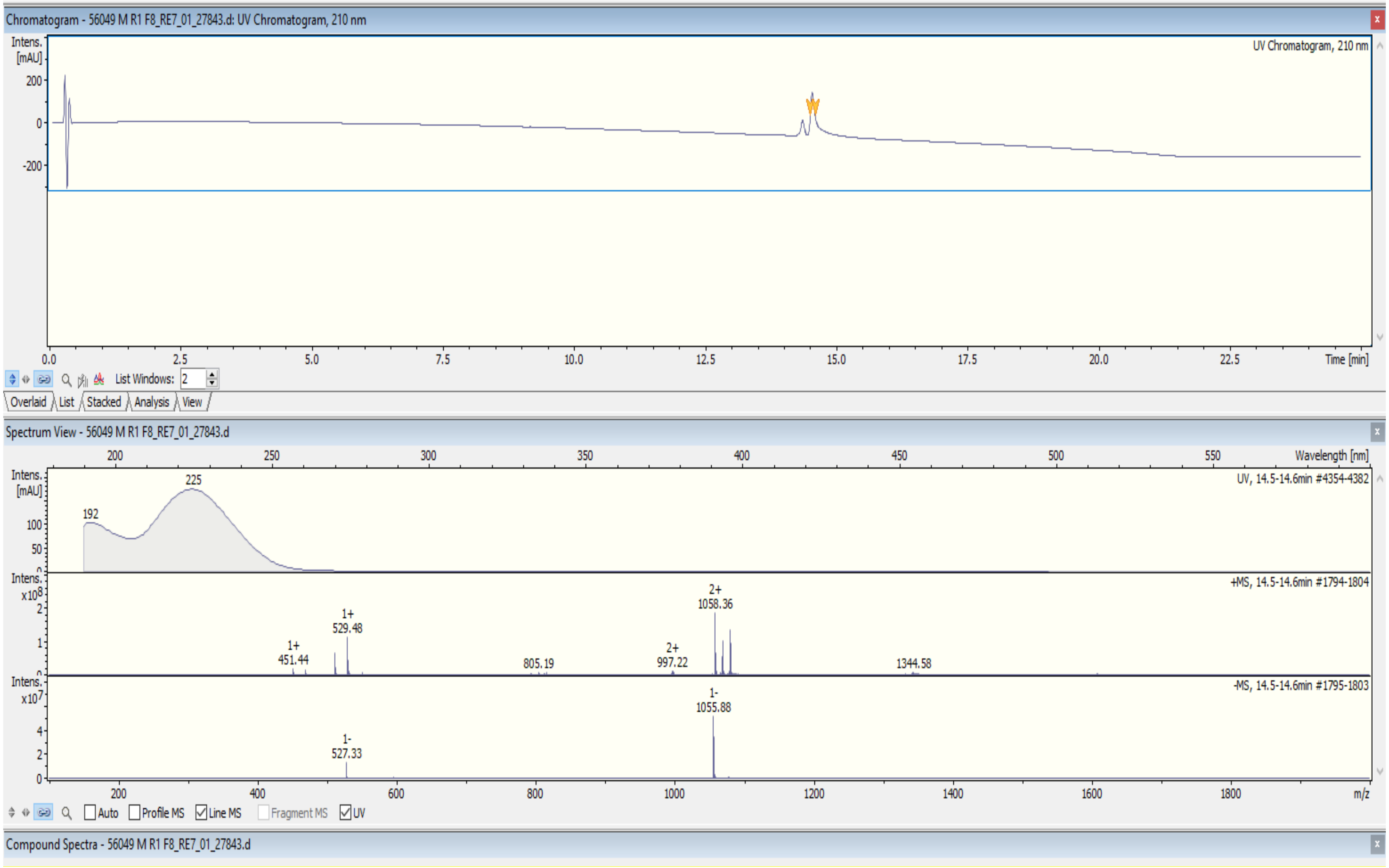


**Figure S26**: ESIMS data for pachymic acid (**12**)

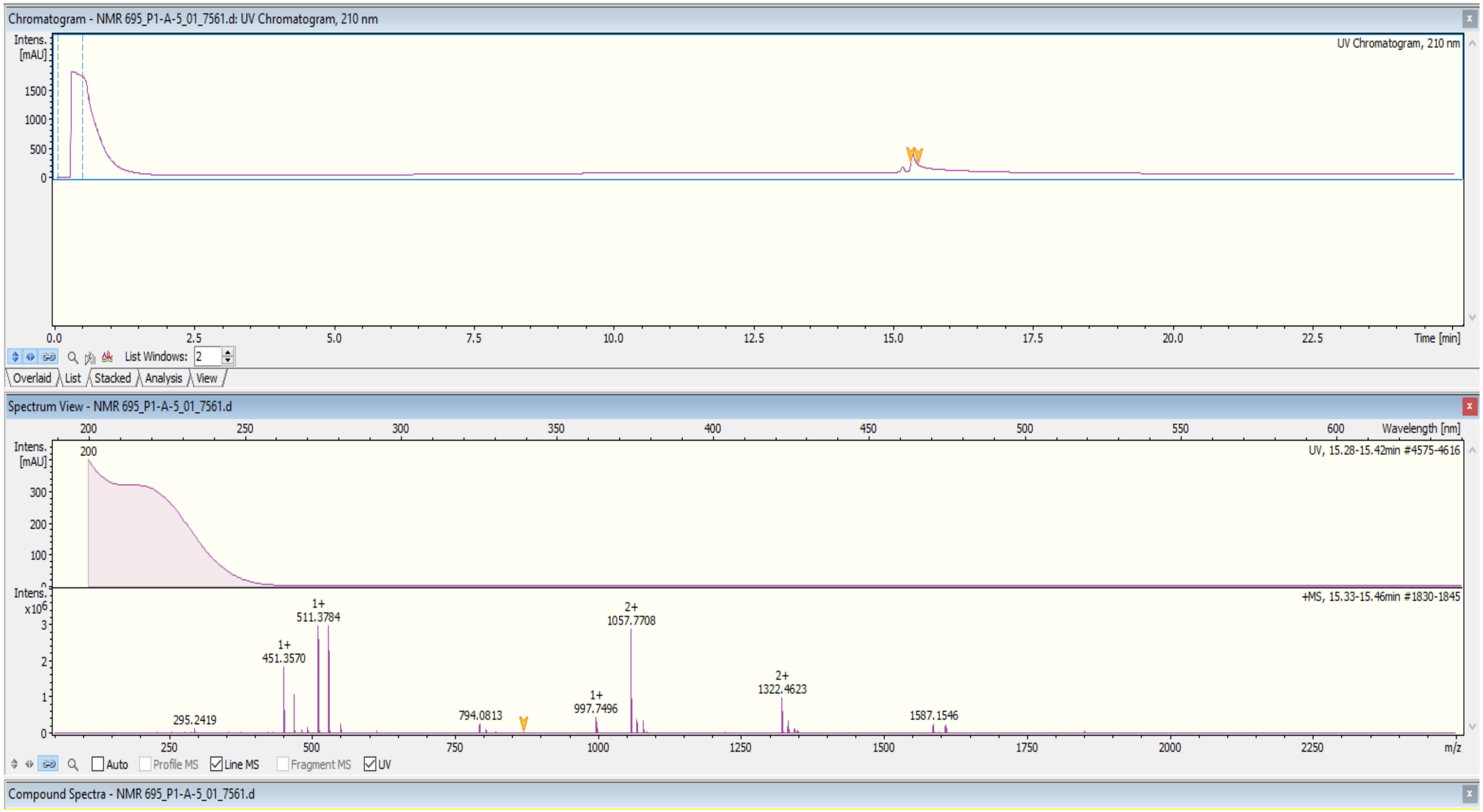


**Figure S27**: HR-ESIMS data for pachymic acid (**12**)

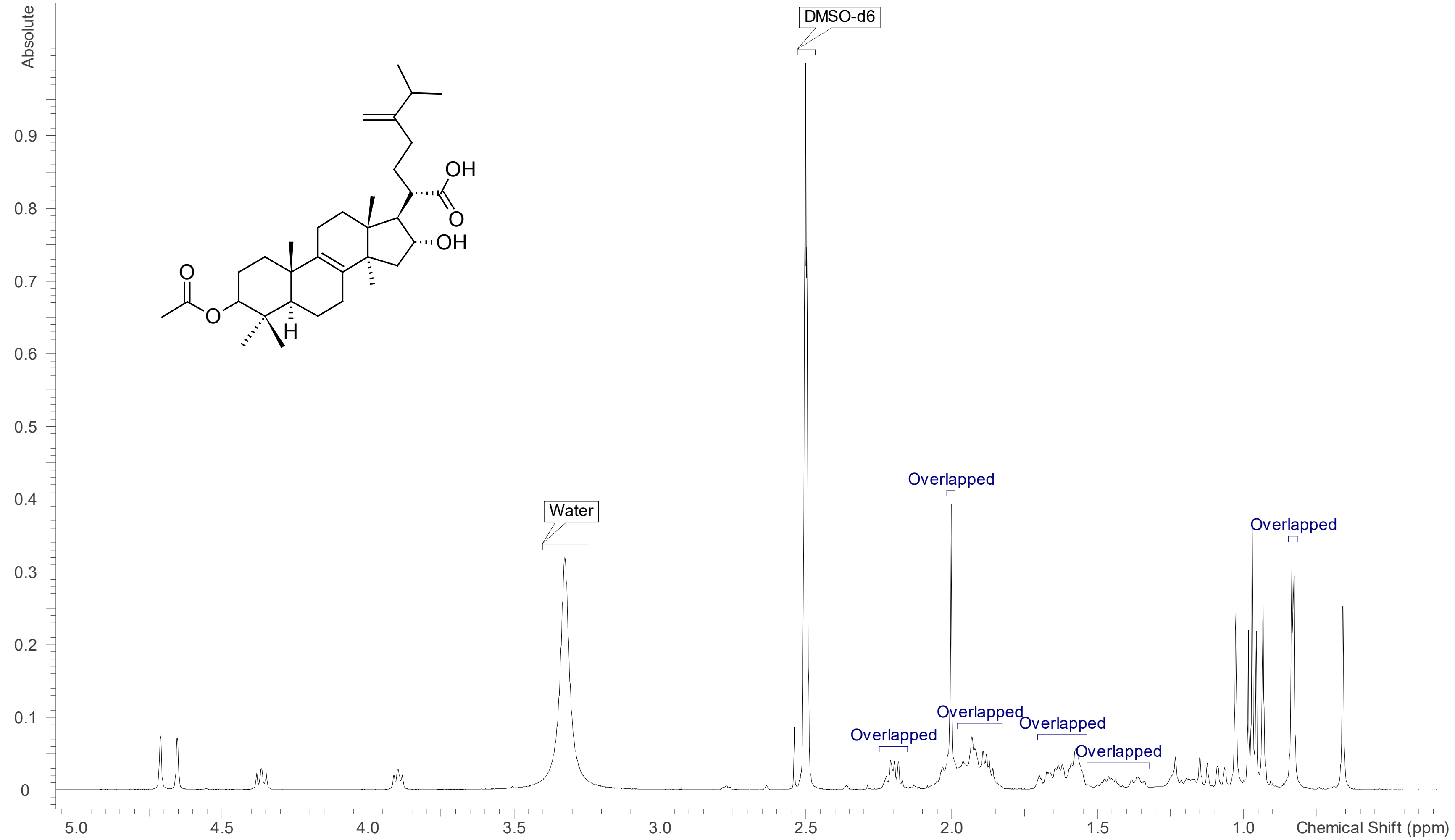


**Figure 28**: 1H NMR spectrum (DMSO- $d_6$, 700 MHz) of pachymic acid (**12**).

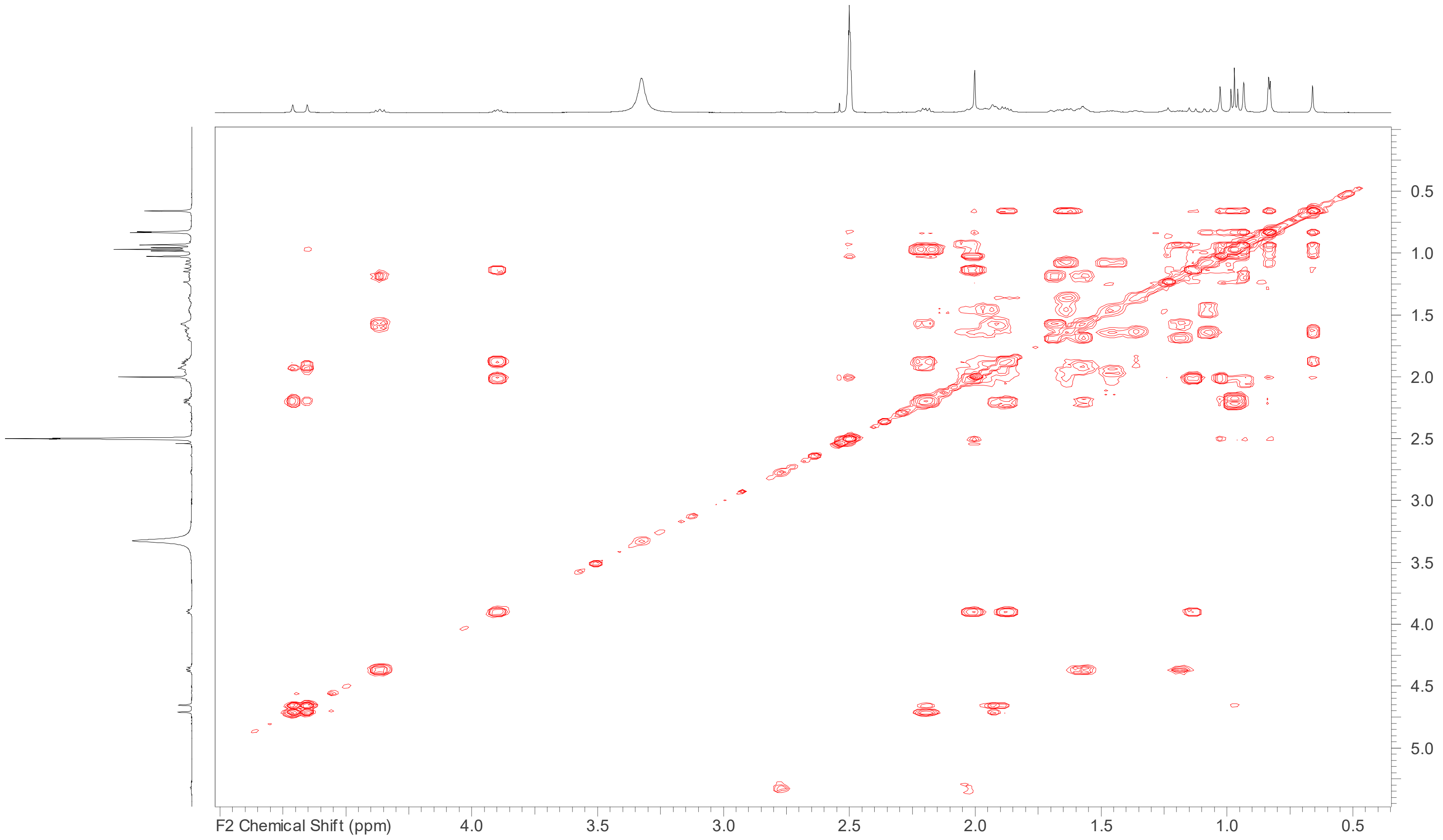


**Figure S29**: COSY spectrum (DMSO-$d_6$, 700 MHz) of pachymic acid (**12**).

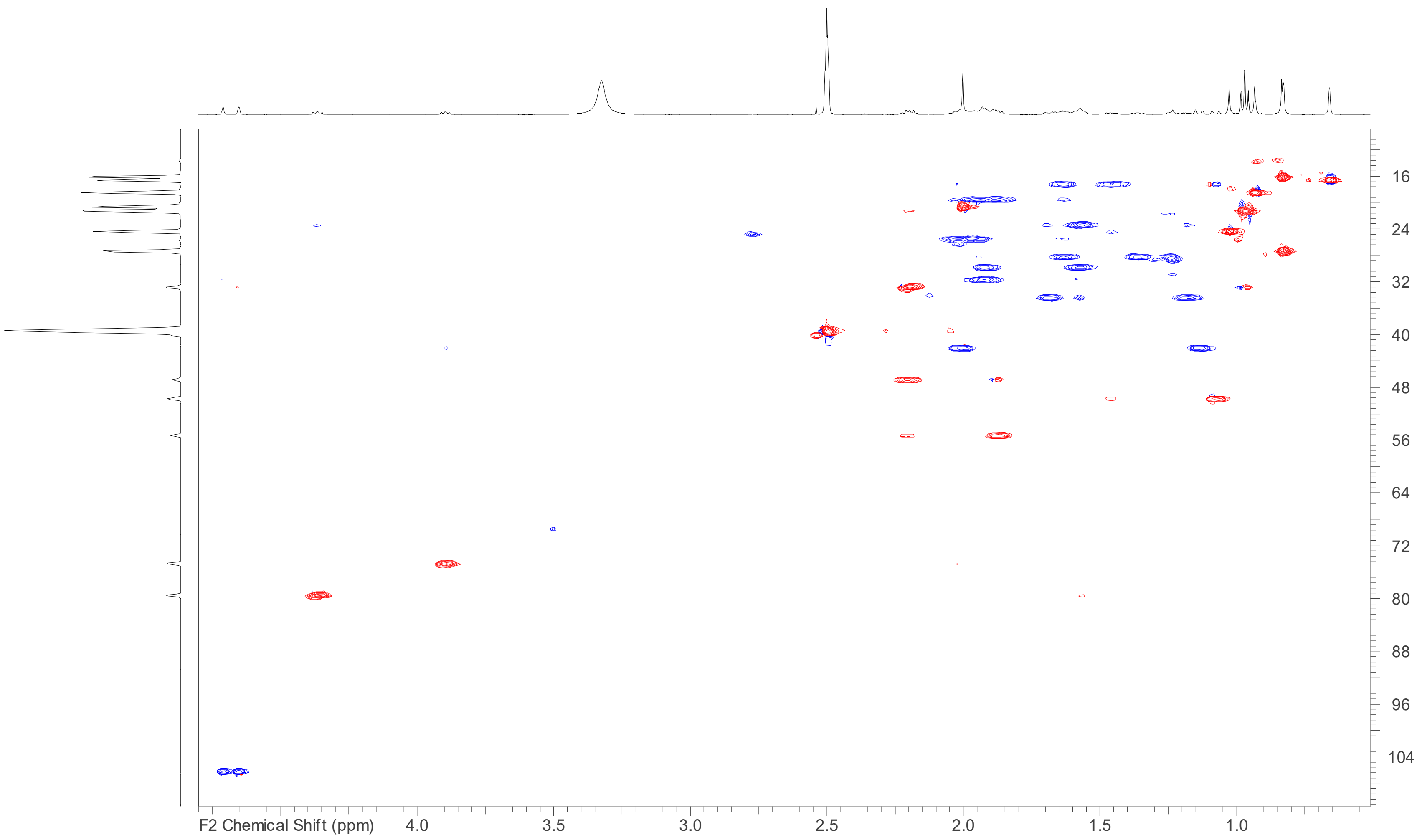


**Figure S30**: HSQC spectrum (DMSO-$d_6$, 700 MHz) of pachymic acid (**12**).

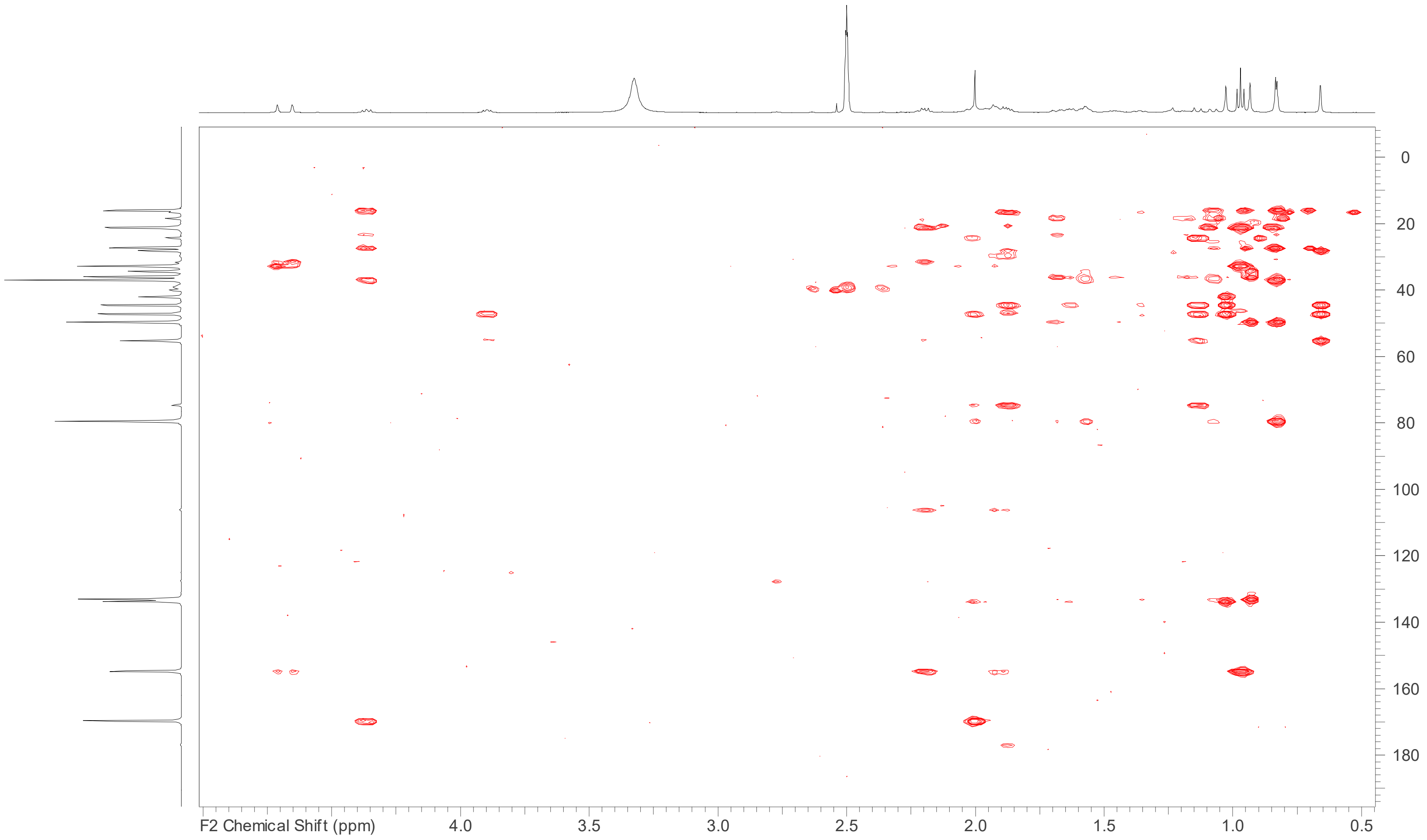


**Figure S31**: HMBC spectrum (DMSO-$d_6$, 700 MHz) of pachymic acid (**12**).

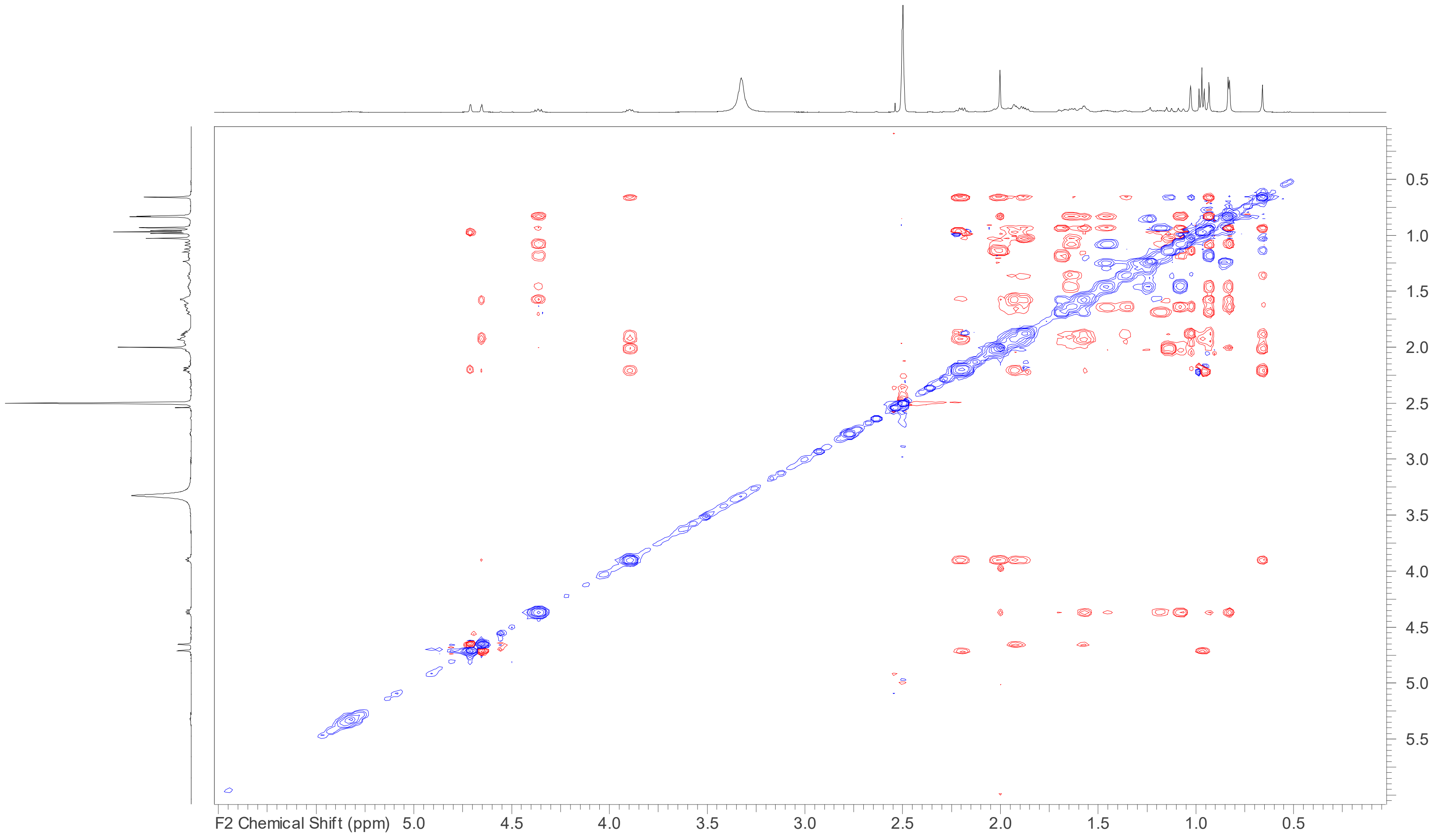


**Figure S32**: ROESY spectrum (DMSO- $d_6$, 700 MHz) of pachymic acid (**12**).

**ITS and LSU sequences of *Antrodia* sp.**

> LSU

GGATTCCCCTAGTAACTGCGAGTGAAGCGGGAAGAGCTCAAATTTAAAATCTGGCGGTCTCTGGCCGTCCGAGTTGTATTCTGGAGAAGTGTTTTCCGTGCTGGA
CCGTGTACAAGTCTCTTGGAACAGAGCGTCATAGAGGGTGAGAATCCCGTCTTTGACACGGACTGCCAGTGCTTTGTGATGCGCTCTCAAAGAGTCGAGTTGTTT
GGGAATGCAGCTCAAAATGGGTGGTAAATTCCATCTAAAGCTAAATATTGGCGAGAGACCGATAGCGAACAAGTACCGTGAGGGAAAGATGAAAAGCACTTTG
GAAAGAGAGTTAAACAGTACGTGAAATTGCTGAAAGGGAAACGCTTGAAGTCAGTCGCGTTGATCAGGACTCAGCCTTGCTTTTGCTTGGTGCATTTTCTGGTTG
ACGGGCCAGCATCGATTTTGACCGTCGGAAAAGGGCTGAGGGAATGTGGCACCTTCGGGTGTGTTATAGCCTTCAGTCACATACGGCGATWGGGATCGAGGAC
CGCAGCACGCCTTTATGGCCGGGGTTCGCCCACGTTCGTGCTTAGGATGCTGGCGTAATGGCTTTAAACGACCCGTCTTGAAACACGGACCAAGGAGTCTAACAT
GCCTGCGAGTGTTTGGGTGGAAAACCCGAGCGCGCAATGAAAGTGAAAAGTTGAGATCCCTGTCATGGGGAGCATCGACGCCCGGGCCTGAACTTCGGTGATG
GTTCTGCGGTGGAGCATGTATGTTGGGACCCGAAAGATGGTGAACTATGCCTGAATAGGGTGAAGCCAGAGGAAACTCTGGTGGAGGCTCGTAGCGATTCTGA
CGTGCAAATCGATCGTCAAATTTGGGTATAGGGGCGAAAGACTAATCGAACCATCTAGTAGCTGGTTCCTGCCGAAGTTTCCCTCAGGATAGCAGAAACTCGTAT
CAGATTTATGTGGTAAAGCGAATGATTAGAGGCCTTGGGGTTGAAACAACCTTAACCTATTCTCAAACTTTAAATATGTAAGAACAACCCGTCACTTGATTGG

> ITS

AGTTCAGCGGGTAATCCTACCTGATCTGAGGTCAAAGGTCAAGATAAATTGTCCTTTAGCAGGACGATTAAGAAGCTGACACCCATACAACATGCTTCACAGAAC
AGTGTAAACAAATTATCACACTGAAGCTGATTCACAAAAGGTTTCAAGCTAATGCATTCAAGAGGAGCTGAACACAGTAGTATCCAGCACACTCCAAATCCAAGC
TCCATTCACAGAAATGAATAGAGTTGAGAATTCCATGACACTCAAACAGGCATGCTCCTCGGAATACCAAGGAGCGCAAGGTGCGTTCAAAGATTCGATGATTCA
CTGAATTCTGCAATTCACATTACTTATCGCATTTCGCTGCGTTCTTCATCGATGCGAGAGCCAAGAGATCCGTTGCTGAAAGTTGTATATAGATGCGTTACACGCA
ATAGACATTCTTTAAACTGAGTTGTGTGTGGGTAAAAACATAGGAAAGACCACAGAGCAAAGTCAATGAAGACTTCACTCCAAGAGCCTAATCTACAGTGTGTGC
ACAGGTGTGAGAGATGGATAATGATCAGGGTGTGCACAATGCCGCAGCCAGCAACAACCCCTTTCAAGATTCATTAATGATCCTTCCGCAGGTTCACCTACGGAA
ACC
